# The genetic architecture of dementia risk: how Alzheimer’s disease vulnerability converges on lipid metabolism and immune cell networks

**DOI:** 10.64898/2026.06.11.731645

**Authors:** Evelina Husén, Zhuoran Cai, Emma Gerrits, Evelina Sjöstedt, Nicholas Mitsios, Tianyu Zheng, Emilio Skarwan, Mathias Uhlén, Åsa Sivertsson, Jan Mulder

## Abstract

Although genome-wide association studies (GWAS) have identified numerous dementia risk loci, their cell-type and tissue-specific contexts remain largely unresolved. We introduced HPA GeneSet Explorer, a statistical pipeline designed to systemically map Genome wide association study (GWAS) disease risk genes across the Human Protein Atlas (HPA). This approach generates a multiscale transcriptomic map of genetic risk, spanning systemic organs, brain regions, and cell types. Applying this framework to Alzheimer’s disease (AD), Lewy body dementia (DLB), and Frontotemporal dementia (FTD) revealed convergent neuronal enrichment in all three diseases. In addition, we identified AD-specific enrichment in immune system and liver associated gene modules, along with DLB-specific enrichment in ciliary modules. By mapping these vulnerabilities in non-demented samples, we provide a blueprint of the baseline vulnerability hotspots that can precede clinical neurodegeneration, offering new targets for disease-specific therapies and biomarker discovery.

## Introduction

Dementia encompasses a spectrum of age-related conditions defined by irreversible progressive cognitive and memory decline. These impairments result from degeneration of neuronal populations and connections that drive cognitive functions. Neurodegenerative disorders are characterized by distinct neuropathological features with overlap in affected brain circuits. This results in overlapping clinical symptoms, complicating diagnosis and identification of disease-specific mechanisms (***Livingston et al., 2024***).

The molecular features associated with these disorders are dominated by disease specific proteinopathies. Alzheimer’s disease (AD) is characterized by aggregated intracellular tangles of hyperphosphorylated tau and extracellular plaques containing *β*-amyloid (A*β*). In contrast, dementia with Lewy bodies (DLB) and Parkinson’s disease (PD) are characterized by aggregates of *α*-synuclein (*α*-syn), whereas frontotemporal dementia (FTD) can be divided in variants with primarily pathological inclusions of tau, TAR DNA-binding protein 43 (TDP-43), or the rarer variant with FUS (fused in sarcoma) (***Bahia et al., 2013***; ***Wilson et al., 2023***). Despite decades of research into these markers and their role in pathogenesis and disease progression, disease-modifying therapies targeting main proteinopathies in dementia fail in preclinical stages or have so far shown very little effect on cognitive decline, as in the case of A*β* targeting antibody therapies (***Espay et al., 2024***). This gap in treatment stems, in part, from a fundamental paradox: while several of these proteins aggregate in disease and degenerate distinct brain circuits, they are also ubiquitously expressed throughout the nervous system and periphery (as shown in the HPA gene pages 1 link 1-4).

Familial and early onset disease variants provide valuable insights into disease etiology. The rare cases of familial early onset dementia are typically linked to autosomal dominant mutations with high penetrance, such as those in *amyloid beta precursor protein* (*APP*), *presenilin 1* (*PSEN1*), and *presenilin 2* (*PSEN2*) for AD, or *synuclein alpa* (*SNCA*) and *leucine rich repeat kinase 2* (*LRRK2*) in DLB and PD, or *C9orf72*, *microtubule associated protein tau* (*MAPT*), and *progranulin* (*GRN*) in FTD. Mechanistically, these mutations drive pathology by either increasing the aggregation propensity of proteins, altering their proteolytic processing, or exhausting the cellular protein quality control system (***Wilson et al., 2023***). However, these familial variants represent only a minor proportion of the total patient population. The majority of dementia cases are sporadic and have a complex genetic and environmental architecture that lacks a single clear causative mutation (***Reitz et al., 2011***; ***Orme et al., 2018***; ***Rohrer et al., 2009***; ***Wirdefeldt et al., 2011***).

To untangle the complex genetic architecture of sporadic cases, Genome-Wide Association Studies (GWAS) have identified hundreds of loci associated with altered disease risk. GWAS are based on genome-wide detection of genetic variants, most commonly single nucleotide polymorphisms (SNPs). These studies are in large populations to link SNPs with disease or disease-related traits. Risk variants are associated with an increased risk of disease, but their causative effects are often low or modest, of low penetrance, and although these mutations increase vulnerability, they require other genetic (polygenic risk) or environmental factors to initiate or progress disease. Nevertheless, these extended lists of disease-associated risk genes provide valuable clues to the disturbed biological pathways and cellular processes that drive pathology. Several GWAS on AD have been published and together with large-scale meta-analyses have provide integrated lists of disease-associated genes. In summary, risk genes associated with AD are enriched for protein coding genes involved in A*β* processing, structural and cytoskeletal proteins, lipid metabolism, oxidative stress and immune responses, particularly microglia (***Bellenguez et al., 2022***; ***Zhang et al., 2023***; ***Kunkle et al., 2019***; ***Sherva et al., 2025***). Alphasynucleinopathies such as PD, Parkinson’s disease dementia, and DLB show a genetic risk enriched in components of protein degradation pathways and lipid associated pathways (***Fanning et al., 2020***; ***Sanghvi et al., 2020***). The few studies investigating genetic risk factors associated with FTD highlight the role of the immune system (HLA locus), lysosomal functions, glucocorticoid signaling, RNA processing, and protein degradation (***Ferrari et al., 2014***; ***Broce et al., 2018***; ***Mishra et al., 2017***; ***Raffaele et al., 2019***). Notably, many risk genes for FTD are also associated with amyotrophic lateral sclerosis (ALS), a neurodegenerative disease that affects motor neurons in the spinal cord and brainstem, suggesting shared pathological processes and selective vulnerability (***Raffaele et al., 2019***). Despite the identification of numerous genetic risk loci, the molecular basis of selective vulnerability and the extent to which these genetic risks converge across different dementias remain unresolved. Characterizing the expression patterns of these risk genes is therefore essential to move beyond statistical associations and toward a systems level mechanistic understanding of neurodegeneration.

To address these challenges, we developed the HPA GeneSet Explorer pipeline, a computational framework designed to map the expression of GWAS-defined risk genes throughout the human body. In this study, we used the pipeline to investigate three diseases that lead to dementia, namely AD, DLB/PD, and FTD/ALS. Using data from the Human Protein Atlas (HPA), our approach utilizes human, non-disease, gene expression profiles to identify vulnerability signatures and their extended molecular network. This strategy bypasses the limitations of disease tissue, where terminal neurodegeneration often obscures the initial drivers of vulnerability. Through this pipeline, we characterize the enrichment of dementia risk genes across biological scales; i) organs and tissues, ii) brain regions, and iii) cell types. The pipeline further organizes risk genes based on enrichment into functional gene modules, again at the mesoscopic levels of body, brain, and cell types. By applying this deconvolution approach in combination with functional enrichment analysis, we identify widespread molecular processes and define those uniquely associated with dementia-susceptible networks and disease-specific signatures describing vulnerabilities and points of clinical intervention.

## Results

To resolve the genetic risk at the level of cell and systemic mechanisms in dementia, we developed the HPA GeneSet Explorer, a statistical framework designed to integrate GWAS-identified risk genes with the HPA. This approach allowed us to map the genetic vulnerability of AD, DLB/PD (hereafter mentioned only as DLB), and FTD/ALS (hereafter mentioned only as FTD) across a multiscale transcriptomic landscape. The HPA provides classification of genes based on co-expression analysis, for a multiscale approach we utilize three HPA datasets. The data encompasses enrichment of genes in tissues and organs, brain regions, and cells as well as gene modules containing genes with shared distribution patterns across biological scales (that are visualized in UMAPs, HPA links 5-7 in 1). The significance of enrichment of gene modules was assessed using a tiered framework of Fisher test significant fold enrichment (*p*_*nominal*_ < 0.05 and false discovery rate (FDR) *p*_*adjusted*_ < 0.1) and Monte Carlo simulations of enrichment (10^6^ permutations, *p*_*empirical*_ < 0.05), ensuring a balance between stringent error control and the sensitivity to capture subtle biological signals.

### Dementia risk genes are often disease-specific, yet their expression patterns are broadly distributed across tissues, brain regions, and cell types

Ontology-based searches in the GWAS Catalog were used for the data retrieval step of the HPA GeneSet Explorer pipeline and resulted in 530 AD, 318 DLB, and 230 FTD risk genes, respectively. After selecting genes of significance *p* < 1·10^−5^ and mapping to the three HPA resources (for details, see Methods), the final datasets included 453 AD, 278 DLB, and 219 FTD risk genes, as shown in panel A 1. Neurodegenerative diseases are known to share several pathological mechanisms, but the risk gene-sets of the investigated dementias appear to be disease-specific, as shown in panel A 1. The limited overlap of risk genes indicates disease-specific genetic architectures and that each disorder is driven by distinct molecular systems. However, two risk genes are shared between AD, DLB and FTD, *major histocompatibility complex, class II, DR alpha* (*HLA-DRA*) and *major histocompatibility complex, class II, DR beta 5* (*HLA-DRB5*). Both genes are enriched in macrophages of the central nervous system (CNS) and are not specific to any brain region (see 1 HPA-links 8 and 9). In general, the investigated dementia risk genes are not selectively expressed in the brain, peripheral tissues, or specific to a certain type of brain cell, but broadly expressed in several tissues and organs, as shown in panel B-D 1 and 1. However, among the small subset of genes that showed enrichment, brain enrichment is the most common in all dementias. Together, these findings reveal that the genetic risk of dementia is shaped by disease-specific molecular networks of widely distributed gene expression.

The HPA tissue, brain region, and single cell data provides classification of genes individually and based on co-expression analysis at the different biological scales. The resulting gene modules are visualized in UMAPs together with annotations of specificity and/or function of the related protein, based on over representation analysis using several biological databases as well as tissue/brain region/ cell type specific information. Of the filtered risk gene-sets, 232 AD, 101 DLB, and 77 FTD risk genes were enriched at *p*_*nominal*_ < 0.05, in a gene module in at least one of the three biological scales (a total of 396 unique genes). For visualization of risk genes in the separate UMAPs see 4,5,and 6. However, 132 AD, 56 DLB, and 34 FTD risk genes were significant at *p*_*adjusted*_ < 0.1. Although the total number of enriched modules varied by disease and resolution, a consistent core of gene module signatures emerged.

**Figure 1.**
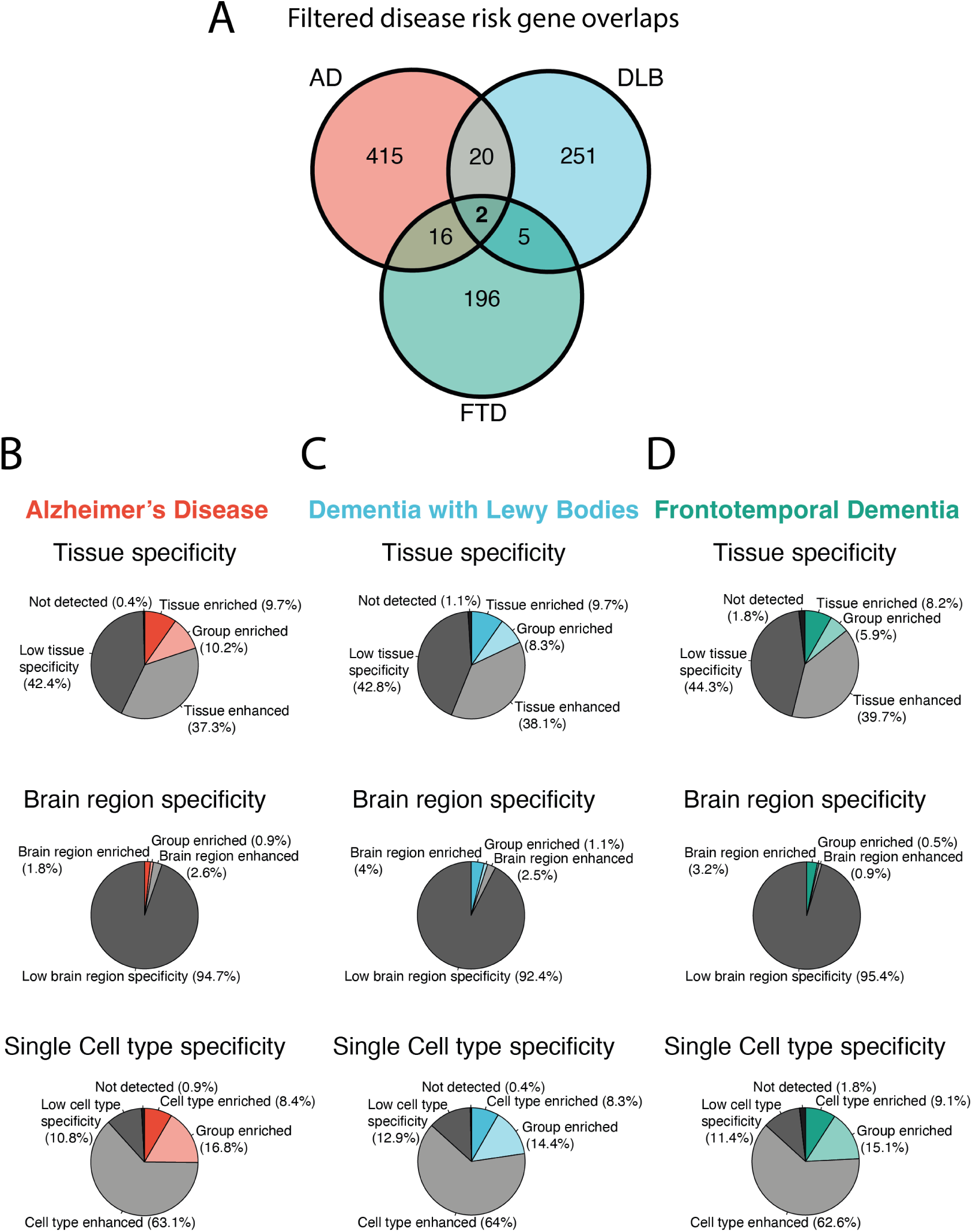
Dementia risk genes exhibited broad systemic expression. A: Venn diagram visualizing the number of filtered risk genes and their overlap across diseases (AD: red, DLB: blue, and FTD: green). B-D: Pie charts depicting the degree of specificity of the neurodegenerative disease risk genes at tissue, brain region, and cell level (again AD: red, DLB: blue, and FTD: green). The specificity categories from highest to lowest specificity are defined as: tissue/brain region/cell type enriched entails ≥ four-fold higher mRNA level in that tissue/brain region/cell type compared to all other tissue/brain region/cell type; group enriched entails ≥ four-fold higher average mRNA level in a group of 2-5 tissues/brain regions or 2-10 cell types compared to all other; tissue enhanced entails ≥ four-fold higher mRNA level in a particular tissue/brain region/cell type compared to average level in all other; and low tissue/brain region/cell type specificity mRNA is detected but the levels are not elevated in any tissue/brain region/ cell type compared to the other. See 1 for the specific compartments of the enriched genes.

Gene modules enriched with disease risk genes, whether nominal, FDR adjusted, and/or empirical significance, were used to define disease-specific gene signatures. Together, the nominal, FDR, and Monte Carlo significant modules paint a consensus picture that converges on a set of biologically coherent signatures that form the basis for subsequent drug target and repurpose analysis. Gene modules will from now on be referred to as modules.

### The neuronal risk signature: dementias share risk gene enrichment in neuronal signaling

In line with neuronal degeneration in dementia, significant enrichment of risk genes from all three dementias was identified in modules associated with neuronal signaling. As shown in panel E-F of 2, 3, and 6, two modules associated to neuronal signaling/synaptic function (cell modules 65 and 34) are enriched in AD, DLB, and FTD. A third neuronal signaling module (module 69) also showed enrichment, but only converging AD and FTD. The neuronal signaling risk signature is further reinforced by enrichment in tissue and brain modules, based on Monte Carlo simulations and nominal fold enrichment, associated with neurons (tissue: module 64 for AD, 61 and 65 for DLB, and 22 FTD; brain region: module 38 for AD and FTD, module 53 for FTD and DLB, and 2 for DLB), as shown in panel A of 2, as Monte Carlo simulation output in 1 and 3, and in the specific UMAP projections in 4 and 6. Taken together, 186 unique dementia risk genes make up the neuronal signaling signature, specifically 82 AD (35% of all enriched AD genes), 62 DLB(61% of all enriched DLB risk genes), and 53 FTD (69% of all FTD risk genes) risk genes respectively. These results indicate a cross-dementia vulnerability in neuronal gene-regulatory processes, particularly in networks involved in synaptic communication and maintenance.

### AD immune risk signature: AD risk genes are enriched in modules associated to immune cells specific of both the periphery and the CNS

In contrast to both DLB and FTD, AD risk genes were enriched in modules specific to peripheral and CNS immune cells. The risk signature is a consensus of both FDR and Monte Carlo enrichment and centered on the AD risk gene specific enrichment. As shown in panel C-F of 2, 1 and 2 AD risk genes are enriched in modules associated with B-cells (cell module 73) and CNS macrophages and microglia (brain module 20). Furthermore, based on Monte Carlo simulations and nominal fold enrichment, the macrophage-associated module (cell module 58) is also part of the signature, as shown in panel E-F of 2 and 3. These enrichment encompasses 45 of the AD risk genes (19% of the module enriched AD genes, *p*_*nominal*_ < 0.05). 7 additional genes (out of the nominally enriched AD risk genes) were identified to be enriched or enhanced in immune cell populations and tissues, although not assigned to immune modules (see 1, HPA links 10-18). AD risk genes are enriched in immune cells and several AD risk genes encode immune system components, no modules associated with immune tissues (such as the thymus) are enriched by AD risk genes. Together, these data suggest an immune vulnerability signature in AD, that is not brain specific but shared with other tissue resident immune cells.

### AD liver associated risk signature: AD risk genes are enriched in liver associated gene modules and the signature has overlaps with the immune gene signature

Beyond the immune risk signature, an AD liver associated risk gene signature becomes apparent, which is not present in DLB or FTD. As seen in tisssue modules *p*_*nominal*_ fold enrichment and Monte Carlo simulations, shown in panel A-B of 2 and 3, the liver metabolism and liver plasma protein associated modules (tissue modules 83 and 11) are enriched by AD risk genes and empirically stable. There are two other liver associated modules (tissue module 31 and cell module 62) that are enriched by AD risk genes at nominal significance and included in the liver associated signature, but these are not verified by Monte Carlo simulation or FDR corrected fold enrichment. Since there are several separate liver associated modules in the tissue UMAP dataset, there is a risk that this fragmentation dilutes the biological signal thus increasing the power needed for a measurable enrichment. In total 28 unique genes (12% of the module enriched AD risk genes) were part of the liver and hepatocyte modules. Worth noting is that all AD risk genes enriched in the hepatocyte cell module were also part of the liver associated tissue modules, except *scavenger receptor class B member 1* (*SCARB1*). *SCARB1* has enriched or enhanced expression in both hepatocytes and the liver resident macrophages, Kupffer cells, but the gene is not included in any of the liver associated tissue modules (see 1 link 19). Additionally, of all AD risk genes that are part of modules enriched at *p*_*nominal*_ < 0.05, five genes (*apolipoportein E*,*APOE*; *acid phosphatase 2 lysosomal*, *ACP2*; *FAM20C golgi associated secretory pathway kinase*, *FAM20C*; *interleukin 6 receptor*, *IL6R*; and *tumor protein P53 inducible nuclear protein 1*, *TP53INP1*) have enhanced expression in the liver and/or hepatocytes, although none of them are part of liver associated modules (see 1, HPA links 20-24). These five genes are instead part of modules related to vasculature, astrocytes, macrophages and microglia, and sub-cortical (brain module 12, 19, 20, and 38) as well as cell modules of B-cells and macrophages (cell module 73 and 58).

**Figure 2.**
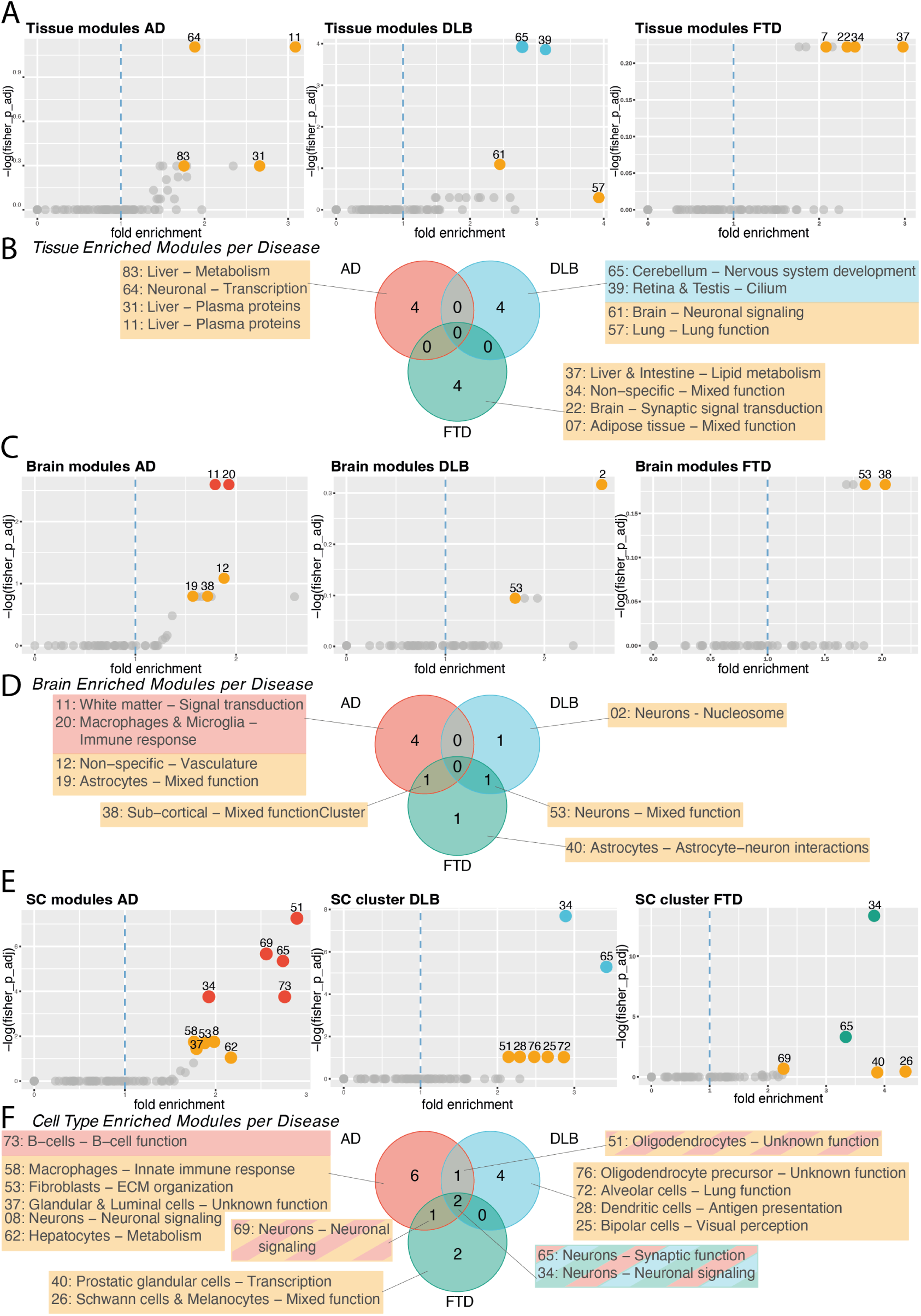
Risk gene fold enrichment across tissue, brain region, and cell gene modules. A and B show tissue module data, C and D brain region mouel data, and E and F cell-type module data. Panels A, C, and E are dot plots depicting the fold enrichment and p-value per module. Each dot represents one module, x-axis is fold enrichment, and y-axis is -log p-value. Dots colored red (AD), blue (DLB), and green (FTD) represent enrichment of *p*_*adjusted*_ < 0.1, while yellow points represent enriched modules of significance *p*_*nominal*_ < 0.05. Panel B,D, and F are venn diagrams illustrating the names and overlaps of enriched modules between disease groups. The numbers is the sum of disease-specific or shared modules, the module numbers and names are summarized in linked boxes. The color coding is as in the dot plots, yellow boxes are significant by <u>*p*_*nominal*_ < 0.05 while red, blue or green boxes are significant by *p*_*adjusted*_ < 0.1. Striped boxes are for modules</u> significant for more than one disease, the red/yellow striped boxes indicate that the module is of significan ce *p*_*adjusted*_ < 0.1 in AD, but *p*_*nominal*_ < 0.05 in DLB or FTD. This indicates the number of interesting gene modules per disease while visualizing similarities and differences across dementias. For enrichment values based on Monte Carlo simulation see 1, 2, and 3

Although the immune and liver associated AD risk signatures seem to define different vulnerability hotspots, they are not entirely isolated. A group of AD risk genes are enriched in modules associated with liver, hepatocytes and immune cells combined, and this kind of overlap is not present between other risk signatures. The group includes genes involved in general metabolic functions including lipid metabolism, such as *apolipoprotein C2* (*APOC2*), *ATP binding cassette subfamily A member 1* (*ABCA1*), *apolipoprotein C1*(*APOC1*), and *lipase C, hepatic type* (*LIPC*). The gene ontology terms of both risk signatures are centered on triglycerides and lipoproteins. However, when analyzing the expression of these themes within brain cells (according to spatial transcriptomics of frontal cortex, 1 link 25) the immune signature genes are mainly expressed by microglia, while the liver associated signature genes are low abundant and not restricted to a single brain cell type (see 7). The overlap between the signatures partially includes genes expressed by tissue resident immune cells in the liver (Kupffer cells) and brain (microglia) genes. When exploring liver associated module enrichment, 19 AD risk genes were identified with expression in Kupffer cells in the periphery and in CNS macrophages and microglia. These include genes involved in functions related to lipoprotein clearance and triglyceride efflux and indicate that cellular process are involved in the initiation and/or progression of the AD. Although there is overlap, more than half of the liver associated risk signature genes are part of liver associated modules and not immune, such as *protein phosphatase 1 regulatory subunit 3B* (*PPP1R3B*), *NHL repeat containing 3* (*NHLRC3*), *O-6-methylguanine-DNA methyl-transferease* (*MGMT*), and *coagulation factor II, thrombin* (*F2*), indicating that these are specifically involved in other biological processes than immune, more specific to liver function.

### DLB cilia risk signature: DLB risk genes are significantly enriched in gene modules associated to cilia function

In contrast to AD and FTD, DLB risk genes display a mesoscopic risk signature associated with cilia and cell polarity. 17 of the DLB risk genes (17% of the module enriched DLB risk genes, *p*_*nominal*_ < 0.05) are enriched in modules linked to cilia-related processes at both tissue and cell type level. The module associated to cilia of retina and testis (tissue module 39) is consensus enriched and the bipolar cell module (cell module 25) is enriched by DLB risk genes according to Monte Carlo simulations and nominal fold enrichment, as shown in panel A-B,E-F of 2, 1, and 3. The tissue module for cilia is defined by gene ontology terms associated to cilia and visual perception and the bipolar cell module is defined by gene ontology terms associated to membrane, dendrites, and synapses. Even though this module is not annotated as cilia specific, its genes cover conserved cellular processes associated with cilia such as vesicle transport, membrane organization, and regulation of signal transduction. These systems are integral to polarity- and cilia-associated cellular mechanisms. Consequently, these modules were grouped within a broader gene signature associated with cilia, reflecting a shared reliance on ciliary-mediated processes rather than organ-specific annotation. Collectively, these modules constitute a DLB-specific vulnerability signature of ciliary-dependent functions.

### Several drugs target dementia risk gene modules leaving a large pharmaceutical potential, as currently only a small fraction have neurodegenerative indications

To evaluate the translational potential of the identified risk signatures, we investigated the overlap between risk genes and drug targets. From this analysis, 1,777 drugs that target the gene products that make up any of the modules enriched by disease risk genes (*p* < 0.05, all the modules noted in 2 on all the biological scales, tissue/brain/cell) were extracted. The drugs were categorized based on direct targeting of risk genes, targeting of risk associated modules, and drugs used for treatment of neurodegenerative disorders or lipid regulation.

As shown in 3, 35 (2%) of the output drugs are used as treatment and symptom management of neurodegenerative diseases, including drugs such as Lecanemab, Levodopa, and Nicergoline (for detailed informatino see Druggable_proteom_dementia.xlsx). Six neurodegenerative drugs target AD specific risk gene products, five target DLB specific risk genes, and one targets AD and DLB specific risk gene products. In total, three anesthetics target risk genes across the three dementias: Sevoflurane, Halothane, and Desflurane. Their overlapping target profiles specifically include gene products of: *gamma-aminobutyric acid type A receptor subunit gamma3* (*GABRG3*) (AD), *gamma-aminobutyric acid type A receptor subunit gamma2* (*GABRG2*) (DLB), and *ATPase plasma membrane Ca2+ transporting 2* (*ATP2B2*) (FTD), all of which are part of the respective neuronal risk gene signature. The fact that these three drugs target the same three risk genes indicates that these protein families are central in the anesthetic effect of the drugs but also further cements the mutual neuronal risk gene signature. Moreover, the output drug list also contains several drugs with repurposing potential and some that have been investigated for neurodegenerative diseases. Such as the malaria drugs Amodiaquine and Hydroxychloroquine, which both target gene products of modules in the immune risk signature, pass the blood brain barrier, and have been investigated in AD (***Matošević et al., 2024***; ***Varma et al., 2023***). Another example is the glucagon-like peptide 1 (GLP-1) and gastric inhibitory polypeptide (GIP) receptor agonist Tirzepatide which has been implicated as a treatment in AD (***Alshehri et al., 2025***). Although our findings associated Tirzepatide to the cilia risk signature so it may have a more targeted potential in DLB and PD (***Yang et al., 2024***; ***Delvadia et al., 2025***). Additionally, lipid lowering drugs were specifically observed in this analysis, due to lipid related immune and liver associated AD risk signatures, and 40 drugs (2.2%) were found targeting dementia and risk signature modules. These include classical drugs such as different statins which have previously been found to lower the risk of dementia (***Filho et al., 2025***). However, several drugs in this list have not been directly associated with dementia treatment. Our intersectional analysis reveals a substantial reservoir of approved and experimental compounds targeting these risk-enriched networks. These findings suggest that the genetic drivers behind the risk or vulnerability to dementia may be more pharmacologically accessible than previously recognized, revealing numerous candidates for drug repurposing.

## Discussion

Identifying the initial molecular drivers of neurodegeneration remains a fundamental challenge, as there are years between the onset of disease and the first manifestation of clinical symptoms. Although genetic risk for dementia generally is distributed across broadly expressed genes, our findings demonstrate that disease-specific vulnerability emerges from selective convergence within molecular and cellular networks. This architecture suggests that vulnerability for neurodegerative disease is determined at the level of biological system dynamics, molecular interactions, and environmental factors. Through our systems approach integrating genetic risk with multiple-scale transcriptomic data, in the HPA GeneSet Explorer pipeline, enrichment was examined across increasing levels of biological organization, from individual genes and tissues, brain regions, and cells to gene networks. This multi-scale approach uncovered robust genetic hotspots that determine vulnerability to dementia.

**Figure 3.**
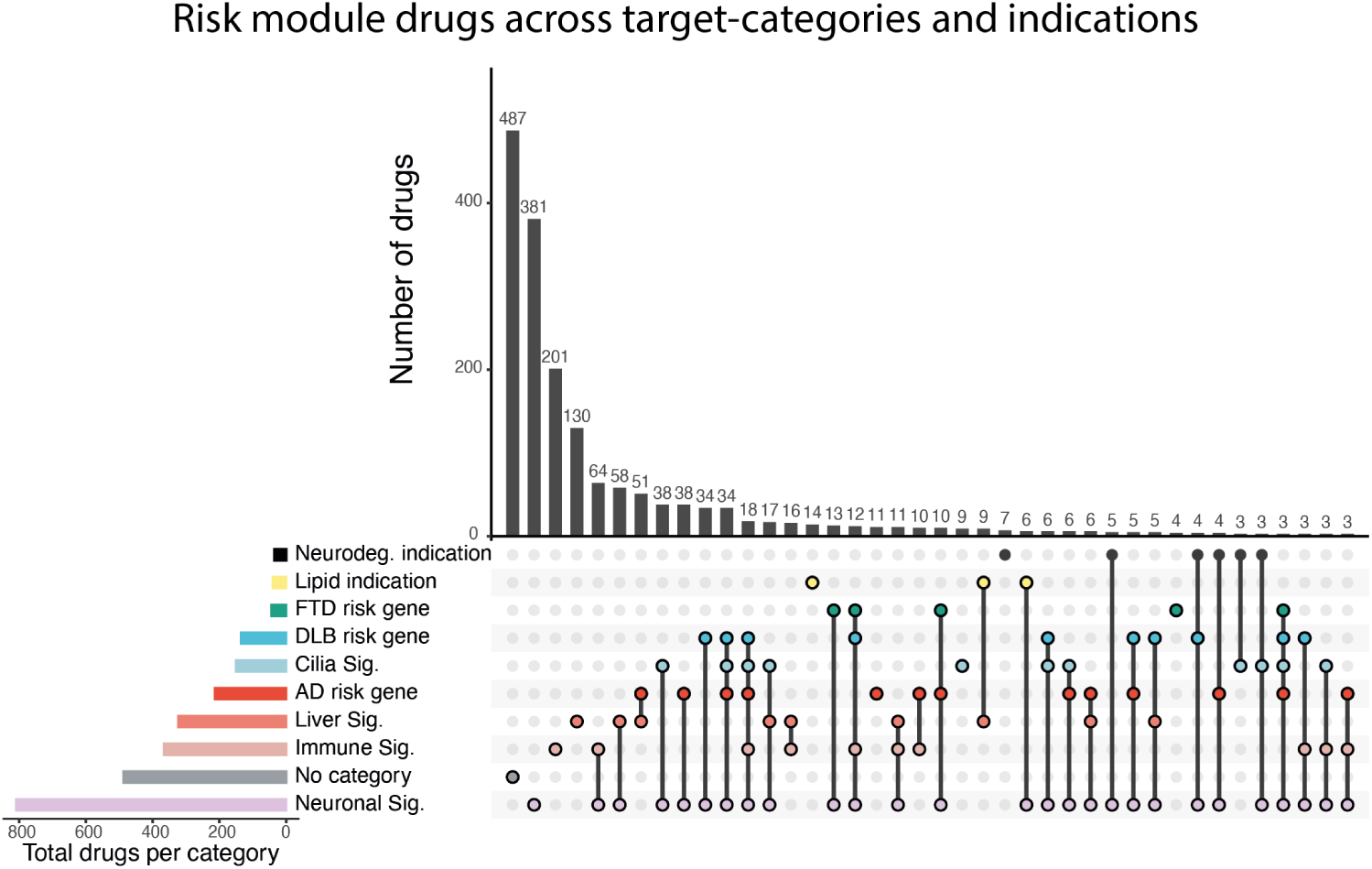
UpSet plot of druggable gene targets across disease risk genes and risk signatures. The drugs were categorized according to whether they have indications specific for neurodegneration or lowering lipid levels, they target gene products of the dementia risk genes, target genes part of any module of the immune/liver/cilia/neuronal associated risk signatures, and part of any gene module (not part of a risk signature) enriched by dementia risk genes (called no category). Each set of horizontal bars represents a category of druggable gene products: disease risk genes (AD, DLB, and FTD), specific drug indications (neurodegenerative disease or lipid lowering), any gene part of risk signature modules (neuronal, immune, liver, and cilia), or no assigned category. Horizontal bars indicate the total number of unique drugs per category. Vertical bars represent the size of each intersection, with dots and connecting lines below indicating which categories overlap. Only combinations specific to > 2 drugs are shown in the plot. A total of 1777 drugs were analyzed, of which only 35 have indications for neurodegnerative disease, and 40 a lipid-related indication. 809 drugs target gene products of genes part of the neuronal risk signature, and the largest drug target overlap is between the neuronal and immune risk signatures (*n* = 64), followed by neuronal and liver associated risk gene signatures (*n* = 58) and AD risk genes and the liver associated gene signature (*n* = 51). Beyond the currently indicated treatments for neurodegeneration we identified an extensive repertoire of experimental and approved compounds with targets mapping directly to the resolved risk signatures, suggesting significant repurposing potential for dementias.

Within this framework, shared neuronal vulnerability emerged in dementias despite a substantial distinction between disease risk genes. Most dementia risk genes are specific to a single disease, yet they are also broadly expressed across the body, implicating an involvement in core cellular processes rather than tissue restricted functions. Despite this wide expression, enrichment analysis revealed convergence of risk genes within neuronal and synaptic modules across dementias. The three diseases share this signature, that is the central hotspot for FTD risk and may be due to FTD and ALS having fewer GWAS based risk genes since they are more rare than both DLB and AD. DLB vulnerability is neuronal intrinsic and not only centered on neuronal signaling but also neuronal cell function through the cilia risk signature. In contrast, AD vulnerability is defined by multi-system risk hotspots, ranging from neuronal signaling to immune cells and lipid metabolism. We show that patterns of enrichment are weak at the level of individual genes or anatomical compartments but become pronounced within modules. Importantly, while the specific enriched modules of the neuronal signature are largely distinct between the diseases, they reflect common functional themes related to neuronal signaling and synaptic function, indicating that shared pathological states arise through disruption of related neuronal systems rather than identical molecular networks. However, these shared genetic networks do not profile specific disease vulnerability, but possibly neurodegenerative disease vulnerability or impaired disease compensatory systems.

Beyond the shared neuronal signature, DLB exhibited a neuronal function-centric genetic vulnerability pattern, with risk genes converging on modules related to cilia and neuronal function. This finding aligns with prior work implicating perturbations of cilia in synucleinopathies. For example, (***Schmidt et al., 2022***) reported a PD-specific effect on cilia in human induced pluripotent stem cell-derived neuronal progenitor cells from sporadic PD patients. Identifying alterations in genes associated with primary cilia function and a shortening of primary cilia length as early cellular alterations associated to mitochondrial dysfunction and disrupted Sonic Hedgehog signaling (***Schmidt et al., 2022***). These results posit that the cilia-associated risk genes are causative vulnerabilities rather than secondary effects. Such alterations are further corroborated by other studies and shown in postmortem tissue of sporadic and familial *LRRK2* PD in striatum (***Khan et al., 2024***; ***Tian et al., 2024***). Consistent with this, *α*-syn has been shown to interfere with ciliogenesis and centrosome function, suggesting convergence on related cellular systems (***Iqbal et al., 2020***). Notably, the rod and cone photo receptor cells of the retina contain large sensory cilia vital for physiological function, and *α*-syn pathology has also been observed in the retina and optic nerve of postmortem tissue of patients with alphasynucleinopathies (***de Ruyter et al., 2023***; ***Wensel et al., 2021***). DLB risk genes uniquely converge on gene networks associated with cilia function and cell polarity, distinguishing DLB from the other dementias examined. While alterations in cilia-related systems have been reported in other neurodegenerative diseases, only DLB risk genes showed significant and consistent enrichment within cilia-associated networks in our analysis, highlighting these pathways as a disease-selective axis of primary vulnerability in DLB (***Serpieri et al., 2025***).

Most strikingly, AD was characterized by the robust multi-system risk signatures, the shared associated with neuronal signaling and the distinct signatures associated with liver and the immune system. The liver and immune system associated hotspots overlap largely in the brain module of macrophage and microglia (module 20), where several of the risk genes are enriched in Kupffer cells and/or hepatocytes, implying a coordinated vulnerability architecture. There is growing observational evidence linking liver function to AD onset and progression. Non-alcoholic fatty liver disease (NAFLD) correlates with a higher risk of cognitive dysfunction and dementia, in particular AD (***Filipović et al., 2018***; ***Weinstein et al., 2022***; ***Lu et al., 2024***; ***Estrada et al., 2019***). In a longitudinal study, Yifei Lu et al found that mid-life NAFLD increases the risk of dementia, while NAFLD later in life did not (***Lu et al., 2024***). An interesting cellular parallel between NAFLD and AD pathology lies in the progressive loss or noradrenergic integrity. In periphery, NAFLD is characterized by significant denervation of liver noradrenergic fibers (***Adori et al., 2021***). This peripheral loss mirrors the central neurodegeneration observed in the locus coeruleus, the main input of noradrenaline to the CNS, which is one of the earliest sites of tau pathology that undergoes sever noradrenergic neuronal loss in AD (***Braak et al., 2011***; ***Weinshenker, 2018***). But through what mechanisms could NAFLD contribute to AD? On the one hand, the connection can be in sequence, e.g. through peripheral clearance of A*β*. Brubaker et al investigated human blood samples and liver tissue to investigate peripheral A*β* clearance. They proposed that A*β* can be cleared through immune adherence and bound to erythrocytes CR1, which are then transported to the liver where Kupffer cells remove them from circulation (***Brubaker et al., 2017***). However, Kupffer cells have currently not been linked to AD and AD risk genes in humans, but are limited to isolated observations in murine experiments (***Yuan et al., 2024***). Hepatocytes have also been implicated in peripheral A*β* clearance, through the low-density lipoprotein receptor-related protein 1 (***Kanekiyo and Bu, 2014***). Consequently, under conditions of liver dysfunction, the loss of peripheral clearing may generate a systemic bottleneck, sequentially accelerating accumulation of A*β* within the CNS and triggering resident immune cells (***Lue et al., 2001***; ***Bianca et al., 1999***; ***Ard et al., 1996***; ***Doens and Fernández, 2014***). The brain immune system, particularly microglia, has long been implicated as a key player in the pathology and development of AD. In mouse models, several AD risk genes have been found to impact microglia function and single nucleus transcriptomics have shown distinct transcriptomic profiles depending on their proximity to A*β* or tau tangles in human AD samples (***Sudwarts et al., 2022***; ***Wang et al., 2015***; ***Aikawa et al., 2019***; ***Jay et al., 2017***; ***Xie et al., 2005***; ***Gerrits et al., 2021***). Perhaps these immune vulnerabilities in the CNS are exposed by liver dysfunction or even by manifestation of peripheral immune vulnerabilities. On the other hand, the connection can also be parallel, e.g. as both NAFLD and AD are associated with metabolic syndrome and insulin-resistance (***Monte and Wands, 2005***; ***Song et al., 2025***; ***Cheon and Song, 2022***; ***Zarghamravanbakhsh et al., 2021***). In 2005 Suzanne M De La Monte et al coined the term type 3 diabetes for AD (***Monte and Wands, 2005***). The naming originated from similarities shared between AD and diabetes mellitus affected cells and systems. In AD, brain metabolism is altered, including decreased glucose utilization and decreased levels of gene expression of insulin, IGF-I, and IGF-II and their receptors(***Steen et al., 2005***; ***Bano et al., 2023***). Notably, liver insulin-resistance, common in diabetes mellitus or prediabetic state, which also are characterized by systemic inflammation, can induce hepatic lipogenesis which is part of the mechanisms leading to NAFLD (***Zarghamravanbakhsh et al., 2021***; ***Anita et al., 2022***). Interestingly, macrophages are drivers of low-grade inflammation which contributes to metabolic dysfunction, and Kupffer cells are described as a central driver of NAFLD pathogenesis (***Zarghamravanbakhsh et al., 2021***; ***Kardinal and Wachten, 2026***; ***Park et al., 2023***). Hence, diabetes may enhance the genetic vulnerabilities for both the peripheral and CNS immune systems, increasing the risk of both AD and NAFLD progression.

Through our analysis of the module enriched risk genes targeted by drugs, we found several of the current drugs used to treat neurodegenerative disease, but we also present a large group of drugs with unutilized potential in treatment of dementias. As part of this output were previously suggested neurodegenerative disease drug candidates, such as Amodiaquine, Hydroxychloroquine, and Tirezpatide (***Matošević et al., 2024***; ***Varma et al., 2023***; ***Alshehri et al., 2025***). This strengthens the exploratory potential of the non-investigated drugs also part of the output. The disease specific risk signatures can be used to further direct the disease implication of the drug, e.g. Tirzepatide which has been investigated preclinically mainly for AD and dementias as a group but based on our findings linking it to the cilia risk signature suggests that it may offer greater therapeutic benefit in specifically DLB and PD (***Alshehri et al., 2025***). Since the most risk genes of the investigated dementias are not causal but genes of generally low penetrance, their effect is cumulative over time, meaning that they generate cumulative damage and not direct disease. Otherwise it would not take decades to cause neurodegneration. Thus, the treatment of these vulnerabilities would need to occur at an early time point and preferably targeting several of the risk signatures to diminish the cumulative risk of several risk mutations and environmental factors. In 2024, the *Lancet* standing Commission presented 14 modifiable risk factors for dementia including points such as education, hearing loss, metabolic diseases, and LDL cholesterol and how to address them over the life course (***Livingston et al., 2024***). Ultimately, our findings offer a framework for preemptive treatment of neurodegenerative disease. By stratifying patient cohorts based on their specific risk signatures, clinicians can identify and therapeutically target distinct vulnerabilities before clinical symptoms emerge. Given the requirement of early intervention, these strategies will rely on long-term clinical surveillance combined with highly tolerable treatments for prolonged maintenance. Transforming these genetic risk insights into proactive interventions is an important step towards preventing dementia and turning the progression into a manageable chronic condition.

## Methods and Materials

The HPA GeneSet Explorer pipeline is developed in R (version 4.5.2) to combine disease genetics and multi-level transcriptomics to explore the molecular landscape of diseases in the human brain and periphery. Here, AD is investigated and compared to DLB and FTD to differentiate if findings are disease specific or common for neurodegenerative diseases.

The input to the HPA GeneSet Explorer is traits or a disease of interest and the output is a PDF-format report summarizing enriched molecular and cellular compartments and gene modules/networks of risk genes across organs, brain regions, and single-cell types, based on data from the HPA version 24 (1 link 26). Hence, the pipeline can be applied to numerous diseases and gene selections. The HPA transcriptomic data encompass 44 tissue types, 193 brain regions and areas, and 81 cell types of 31 human organs and tissues from control donors, which make up the Tissue, Brain, and Single cell type resources. This multi-resolution and multi-angle classification of gene expression makes the HPA GeneSet Explorer a flexible tool that can be applied in any disease context. The integrative approach of the HPA GeneSet Explorer provides a comprehensive overview that provides a means of identifying hotspots where several risk-genes converge on a single cell-type or processes. This enables molecular dissection of pathological processes, identifying disease initiating, adaptive, and maladaptive processes, and provides an overview of other proteins involved in disease associated cellular processes. The work flow is composed of 2 parts, as shown in 4 the initial data retrieval and the analysis and output.

I. Data retrieval: 1) For our analysis specifically, risk genes associated with AD, DLB, and FTD are retrieved from the GWAS Catalog (https://www.ebi.ac.uk/gwas/), using ontology-based search terms. These include Experimental Factor Ontology (EFO), MONDO Disease Ontology, and OBA terms. For each disease, a set of relevant terms are chosen to maximize coverage while avoiding generic or non-specific selections. To enhance our gene discovery for FTD and DLB, ALS and PD were included as terms, respectively, as these diseases have overlapping genetic backgrounds and proteinopathy (the specific search terms are shown in 1). In HPA GeneSet Explorer, all data retrieval is conducted in R using the gwasrapidd package, which allows structured access to the GWAS REST API (***Magno and Maia, 2020***). Gene-to-variant mapping relies on the author-reported genes field provided by the GWAS Catalog, which reflects gene names reported by the original study authors. 2) After the retrieval of gene sets from the GWAS Catalog, the analysis is restricted to variants meeting a suggestive significance threshold (*p* < 1·10^−5^) rather than the conventional genome-wide threshold (*p* < 5·10^−8^), to maximize gene set coverage. This is particularly relevant for diseases where fewer significant loci have been identified throughout the genome, such as DLB and FTD. When a gene is associated with multiple independent loci, only the record with the lowest association p-value is retained to prevent duplicate gene entries. 3) Disease-associated risk genes are matched to HPA gene expression data using HPA custom API queries, non-protein coding genes are implicitly filtered out at this step. For the specific research question of this study, genes not identified at transcript level in the HPA brain datasets were excluded. Moreover, for each gene, metadata is retrieved, including: gene annotation (Gene symbol, Ensembl ID, description, protein class, biological process, molecular function, disease involvement); RNA expression data (tissue/brain region/cell-type specificity and distribution); module assignments (on the HPA the tissue, brain, and single cell transcriptomic data have individually been used to generate gene co-expression UMAPs, where each dot is a gene and the proximity of dots depicts their correlated activity which is used to group them in to clusters/gene modules that represent common biological functions or cell-type specificity); Global expression Tau scores (for tissue, brain, and single-cell); and the API request is configured to retrieve selected columns in uncompressed JSON format for effective parsing.

**Figure 4.**
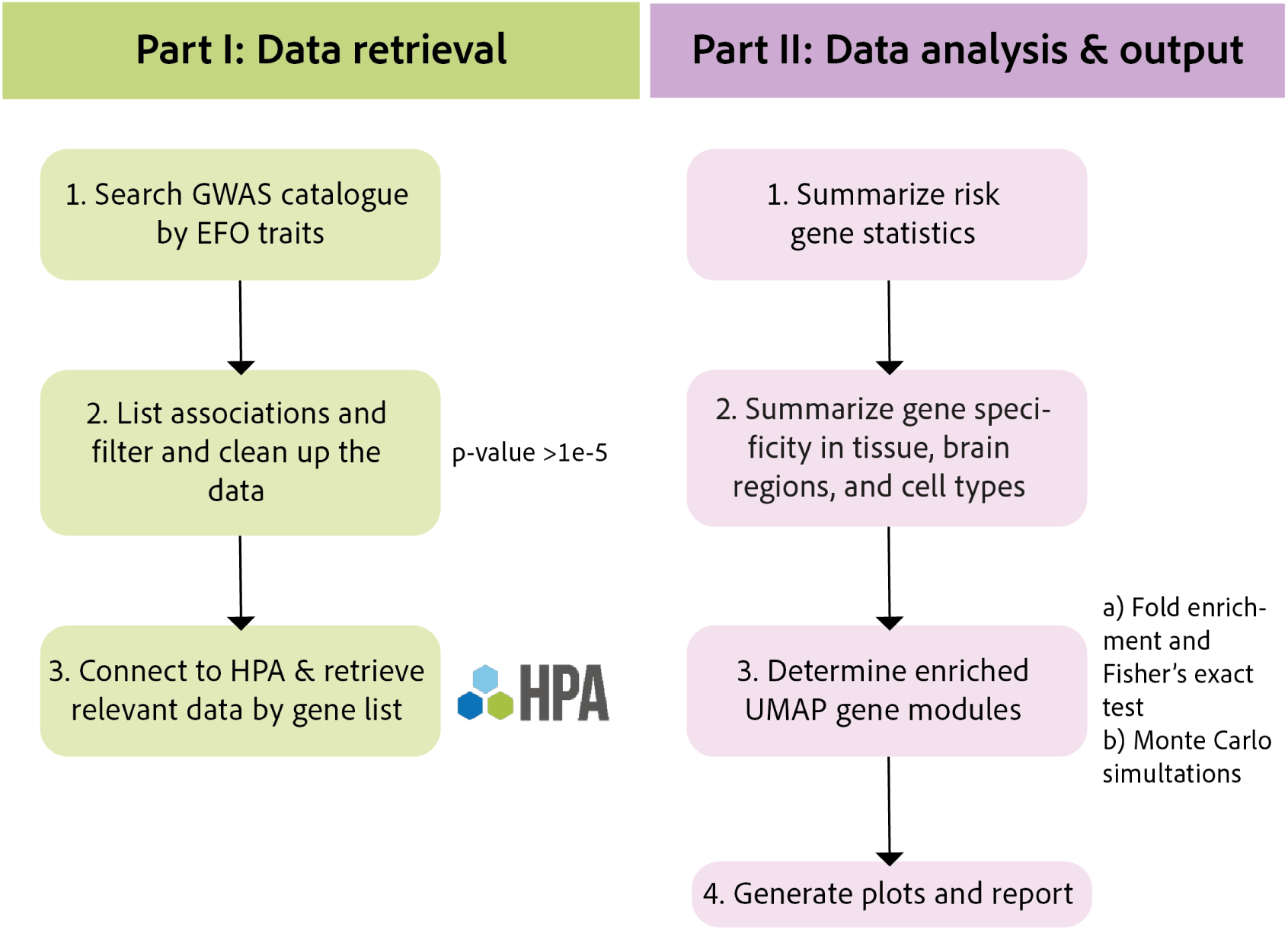
The HPA GeneSet Explorer Workflow. The workflow is divided into I: data retrieval and II: a data analysis component. The data retrieval function searches and retrieves a list of risk genes from the GWAS Catalog using ontology-based terms and filtered by coding status and significance threshold. These genes are then mapped based on expression data for the tissue, brain, and single cell resource of the HPA. In the data analysis component, spatial expression modules are analyzed to identify regions, cell types, and functions enriched for risk genes using statistical tests and assigned to UMAPs of the gene network (see 4, 5, and 6). This approach facilitates spatial interpretation of genetic risk by identifying vulnerable systems and organizing risk genes into discrete functional units. By characterizing the molecular landscape of these disease-associated modules, the investigation pipeline expands beyond individual mutation-based risk genes, providing a broader framework for future analysis of disease-specific vulnerability.

II. Data analysis: 1) the first output is a summary of the risk gene statistics, including the number and fraction of GWAS risk genes retained at each filtering step, as well as the expression and enrichment of individual risk genes across the tissue, brain, and single-cell scales. 2) The HPA provides an overview of gene expression, clustered on their distribution and co-expression across tissues, brain regions and cell-types. This generates UMAPs of modules with similar tissue and/or cell expression patterns. In the HPA GeneSet Explorer pipeline, the risk genes are mapped on UMAP projection gene expression modules of the tissue (83 modules), brain (56 modules), and single cell (80 modules), respectively (see 1 links 5-7). This allows mapping of risk genes onto a low-dimensional transcriptomic space to show vulnerability hotspots of disease risk. The final dataset encompasses; module identities, UMAP x- and y-coordinates, module sizes, and inclusion status (whether a gene is successfully matched to a module). 3) The integrated dataset is used for downstream analyzes to group genes into discrete co-expression modules and subsequently identify which functional, spatial, or cellular units are enriched for disease risk genes. To explore the risk genes enrichment and co-expression of the modules, individual enrichment analysis is performed. Two methods are used to define enrichment that together define a consensus to minimize false positives while recovering biologically relevant findings, method a) fold enrichment and b) Monte Carlo simulations. a) to assess whether disease-associated risk genes are non-randomly enriched in modules, over representation analysis through Fisher’s exact test is used. For each module, the number of observed overlapping genes is computed, defined as the number of disease risk genes present within that module. This is compared to the expected number of overlapping genes, which represents the number of risk genes that are found in a module if they were randomly distributed across all genes in the dataset. The expected overlap is calculated as:

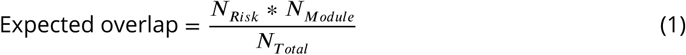

Where N_*Risk*_ is the total number of risk genes retained after filtering, N_*Module*_ is the number of genes assigned to the gene expression module, and N_*Total*_ is the total number of genes in the complete dataset. Then the degree of enrichment is quantified using two methods a) Fold Enrichment (FE):

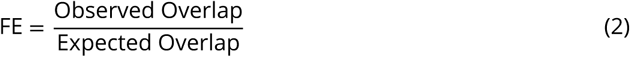

Interpretation of FE values is as follows: *F E* > 1 indicates over-representation of risk genes in the enriched module, *F E* = 1 implies no enrichment beyond random expectation, and *F E* < 1 suggests under-representation (depletion) of risk genes. Statistical significance for each module is determined using a one-tailed Fisher’s exact test (alternative = "greater"), testing specifically for over-representation of risk genes within each module, comparing the number of overlapping versus non-overlapping genes within the module against the background. Modules with a significantly enriched group of risk genes are prioritized for further analysis. Multiple testing correction was applied to the Fisher’s exact test p-values using the Benjamini−Hochberg procedure, applied independently for each biological scale (tissue, brain region, and single-cell type). Modules meeting *p*_adjusted_ < 0.1 were considered FDR-significant. b) Monte Carlo simulations are also used to asses whether disease risk genes are non-randomly enriched within modules. An empirical null distribution is using 1,000,000 iterations, where risk gene assignments were simulated across the complete background gene set using weighted multinominal sampling based on module size. This framework allows identification of high-confidence enriched modules through an empirical p-value (*p*_*empirical*_ < 0.05) and the calculation of Z-scores to quantify the magnitude of deviation from the null distribution. These two approaches enable the identification of hotspot modules that are relevant to the molecular or cellular pathogenesis of the investigated disease or traits, here neurodegenerative diseases. 4) Finally, all output is summarized into a report in a PDF report.

Following the main analysis pipeline, genes from the dementia risk gene-enriched modules were mapped to the DrugBank, where their protein products were queried for approved and experimental drug targets.

In this study the module enriched risk genes for each neurodegenerative disease were investigated in detail. Risk gene signatures were defined based on their expression described integrated across the HPA tissue, brain, and single cell resource. To optimize the balance between sample depth and cell-type taxonomy, our primary analysis was built using HPA version 24. While the subsequent HPA version 25 offers increased resolution that benefits highly granular systems mapping, version 24 provides the ideal equilibrium for our application. Because HPA archives historical versions, users can select their optimal version and our tool is compatible with both version 24 and 25. Moving forward, incorporating user-defined resolution controls within the HPA-platform would further facilitate tailored applications.

## Code availability

All scripts required to reproduce the analyses presented in this study are publicly available at https://github.com/labrat-222/HPA-GeneSet-Explorer. The repository includes modules for gene set retrieval, HPA integration, enrichment analysis, visualization, and automated report generation.

## Data availability

All data used in this study are derived from publicly available resources: the Human Protein Atlas (https://www.proteinatlas.org/) and DrugBank (https://go.drugbank.com/) (***Uhlén et al., 2015***; ***Sjöstedt et al., 2020***; ***Karlsson et al., 2021***; ***Knox et al., 2024***). The following datasets generated during this analysis are available as supplementary files: the full annotated enriched risk gene datasets per disease (Dementia_risk_gene_annotation.xlsx) and the drug and gene product target dataset (Druggable_proteom_dementia.xlsx).

## Appendix 1

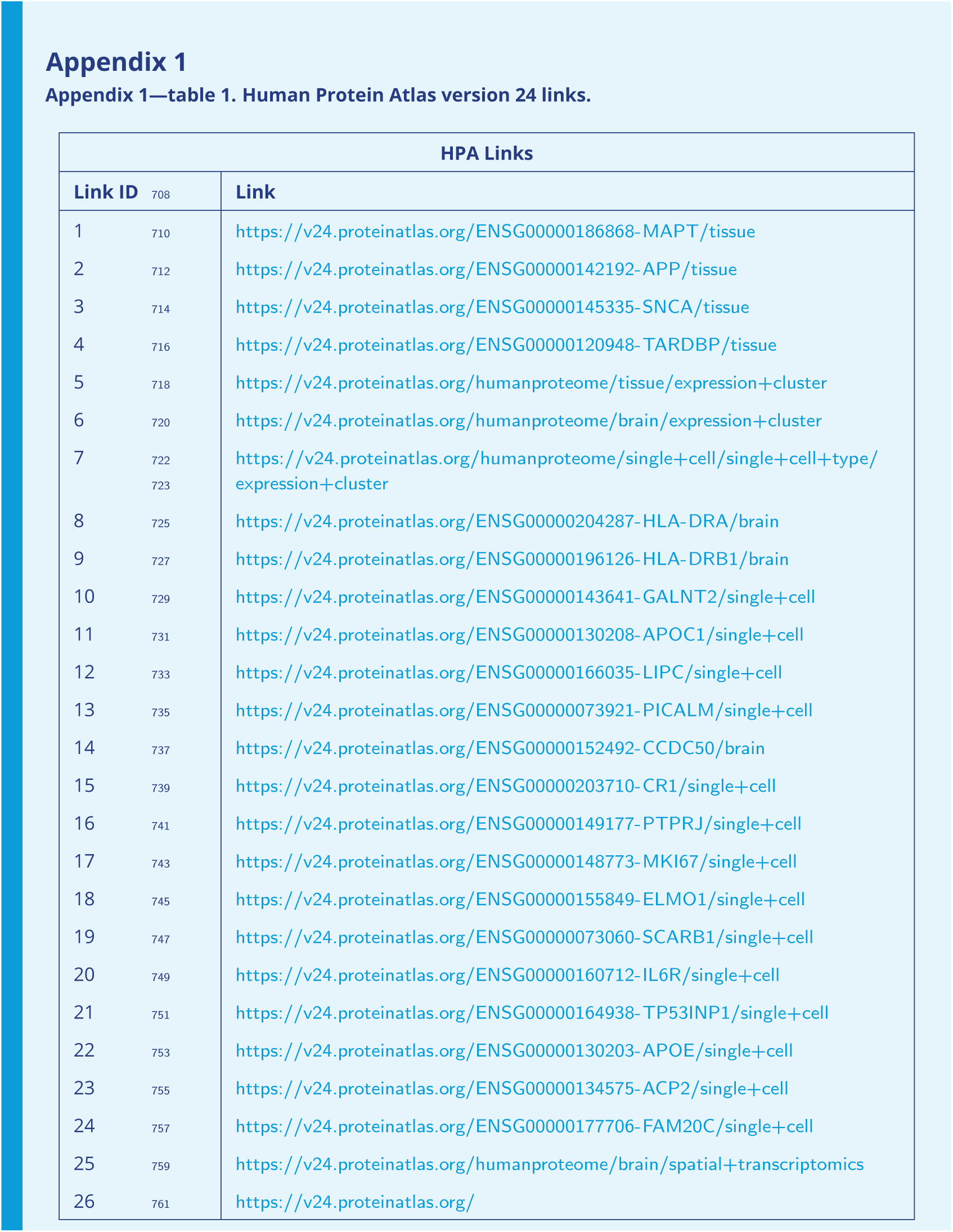

| HPA Links |  |  |
| --- | --- | --- |
| Link ID |  | Link |
| 1 | 710 | https://v24.proteinatlas.org/ENSG00000186868-MAPT/tissue |
| 2 | 712 | https://v24.proteinatlas.org/ENSG00000142192-APP/tissue |
| 3 | 714 | https://v24.proteinatlas.org/ENSG00000145335-SNCA/tissue |
| 4 | 716 | https://v24.proteinatlas.org/ENSG00000120948-TARDBP/tissue |
| 5 | 718 | https://v24.proteinatlas.org/humanproteome/tissue/expression+cluster |
| 6 | 720 | https://v24.proteinatlas.org/humanproteome/brain/expression+cluster |
| 7 | 722<br>723 | https://v24.proteinatlas.org/humanproteome/single+cell/single+cell+type/<br>expression+cluster |
| 8 | 725 | https://v24.proteinatlas.org/ENSG00000204287-HLA-DRA/brain |
| 9 | 727 | https://v24.proteinatlas.org/ENSG00000196126-HLA-DRB1/brain |
| 10 | 729 | https://v24.proteinatlas.org/ENSG00000143641-GALNT2/single+cell |
| 11 | 731 | https://v24.proteinatlas.org/ENSG00000130208-APOC1/single+cell |
| 12 | 733 | https://v24.proteinatlas.org/ENSG00000166035-LIPC/single+cell |
| 13 | 735 | https://v24.proteinatlas.org/ENSG00000073921-PICALM/single+cell |
| 14 | 737 | https://v24.proteinatlas.org/ENSG00000152492-CCDC50/brain |
| 15 | 739 | https://v24.proteinatlas.org/ENSG00000203710-CR1/single+cell |
| 16 | 741 | https://v24.proteinatlas.org/ENSG00000149177-PTPRJ/single+cell |
| 17 | 743 | https://v24.proteinatlas.org/ENSG00000148773-MKI67/single+cell |
| 18 | 745 | https://v24.proteinatlas.org/ENSG00000155849-ELMO1/single+cell |
| 19 | 747 | https://v24.proteinatlas.org/ENSG00000073060-SCARB1/single+cell |
| 20 | 749 | https://v24.proteinatlas.org/ENSG00000160712-IL6R/single+cell |
| 21 | 751 | https://v24.proteinatlas.org/ENSG00000164938-TP53INP1/single+cell |
| 22 | 753 | https://v24.proteinatlas.org/ENSG00000130203-APOE/single+cell |
| 23 | 755 | https://v24.proteinatlas.org/ENSG00000134575-ACP2/single+cell |
| 24 | 757 | https://v24.proteinatlas.org/ENSG00000177706-FAM20C/single+cell |
| 25 | 759 | https://v24.proteinatlas.org/humanproteome/brain/spatial+transcriptomics |
| 26 | 761 | https://v24.proteinatlas.org/ |

## Appendix 2

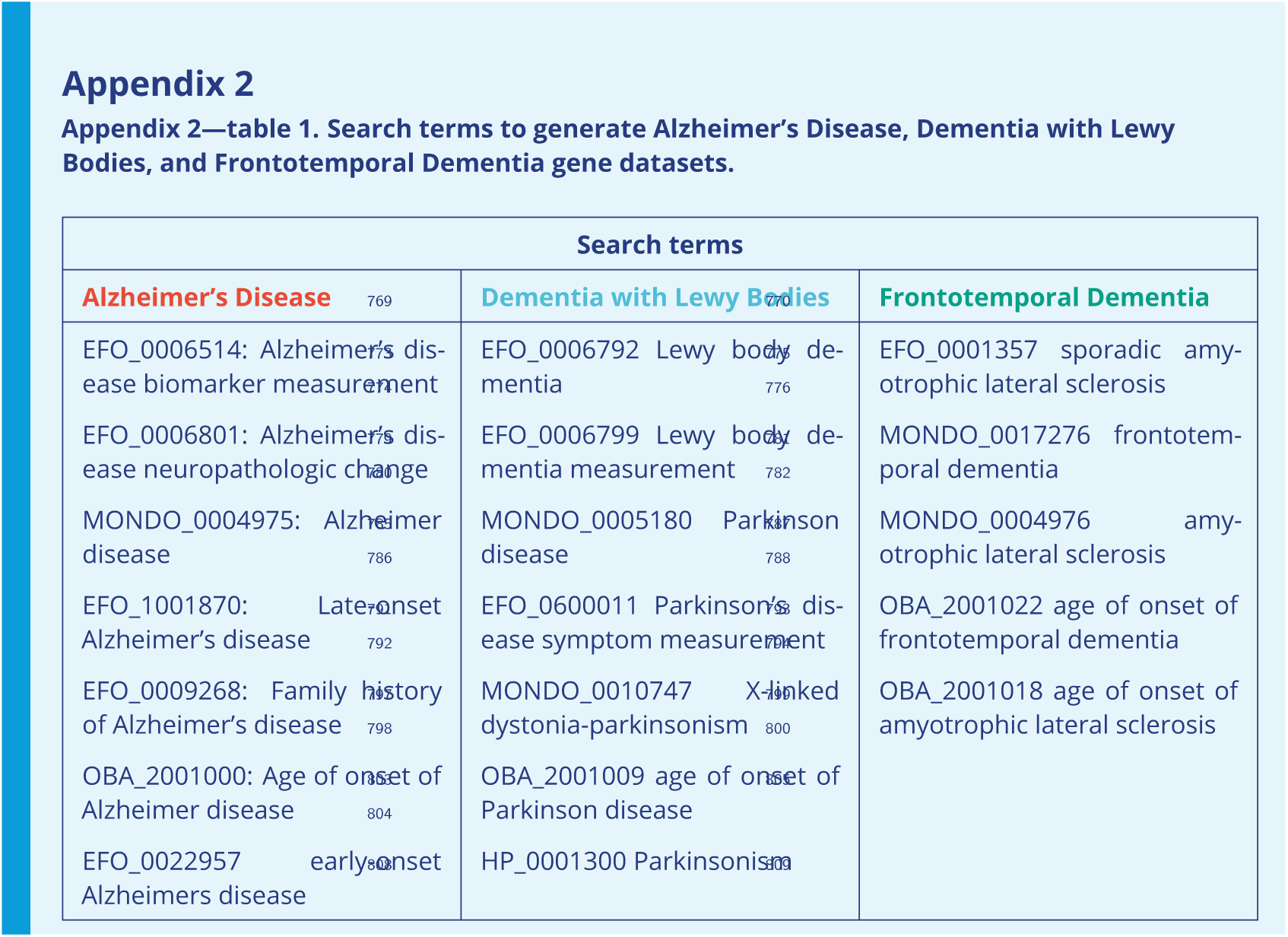

---

**Figure 1—figure supplement 1.**
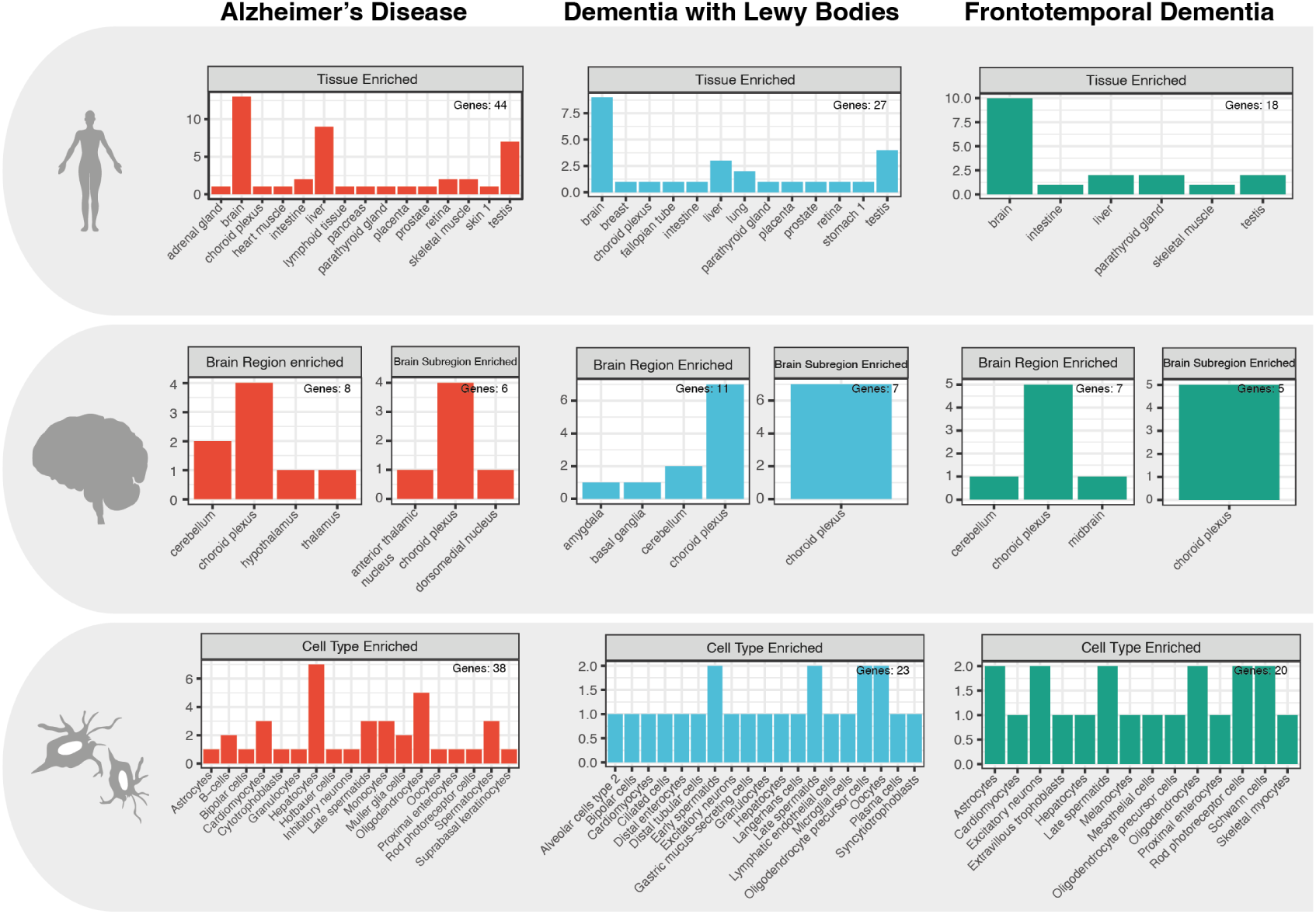
Bar plots of individual risk gene enrichment at the different biological scales. Bar plots representing the risk genes defined as enriched or group enriched in a tissue/brain region/cell type according to the HPA enrichment scoring system mentioned in 1. The y-axis is the number of genes enriched and the x-axis shows the tissue, brain region, or cell type that is enriched. Top panel: tissue, Middle panel: brain regions, and Bottom panel: cell-types. The three diseases are plotted separately and color coded AD: red, DLB: blue, and FTD: green. These plots visualize the low levels of risk gene specific tissue/brain region/cell type enrichment are for each disease.

**Figure 2—figure supplement 1.**
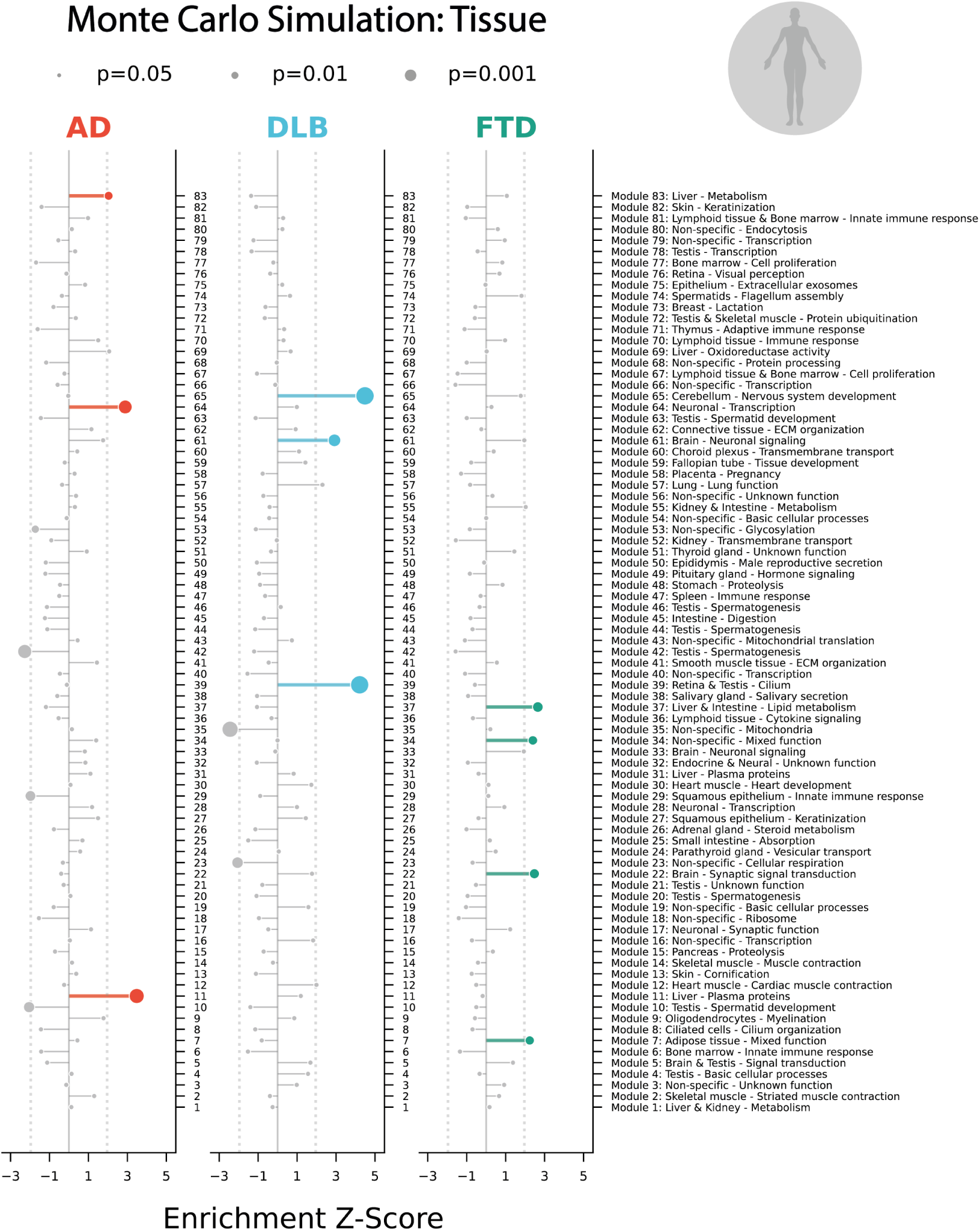
Lollipop plots showing the Monte Carlo simulation disease risk gene enrichment of the tissue gene modules per disease. Enrichment of dementia risk genes in tissue gene modules were analyzed through 10^6^ Monte Carlo simulations. All modules enriched *p*_*empirical*_ < 0.05 are highlighted by color, the gray lollipops are not significant or of negative Z-score (AD: Red, DLB: Blue, and FTD: Green). The size of the plot point indicates the p-value (lower *p*_*empirical*_ has a larger dot size).

**Figure 2—figure supplement 2.**
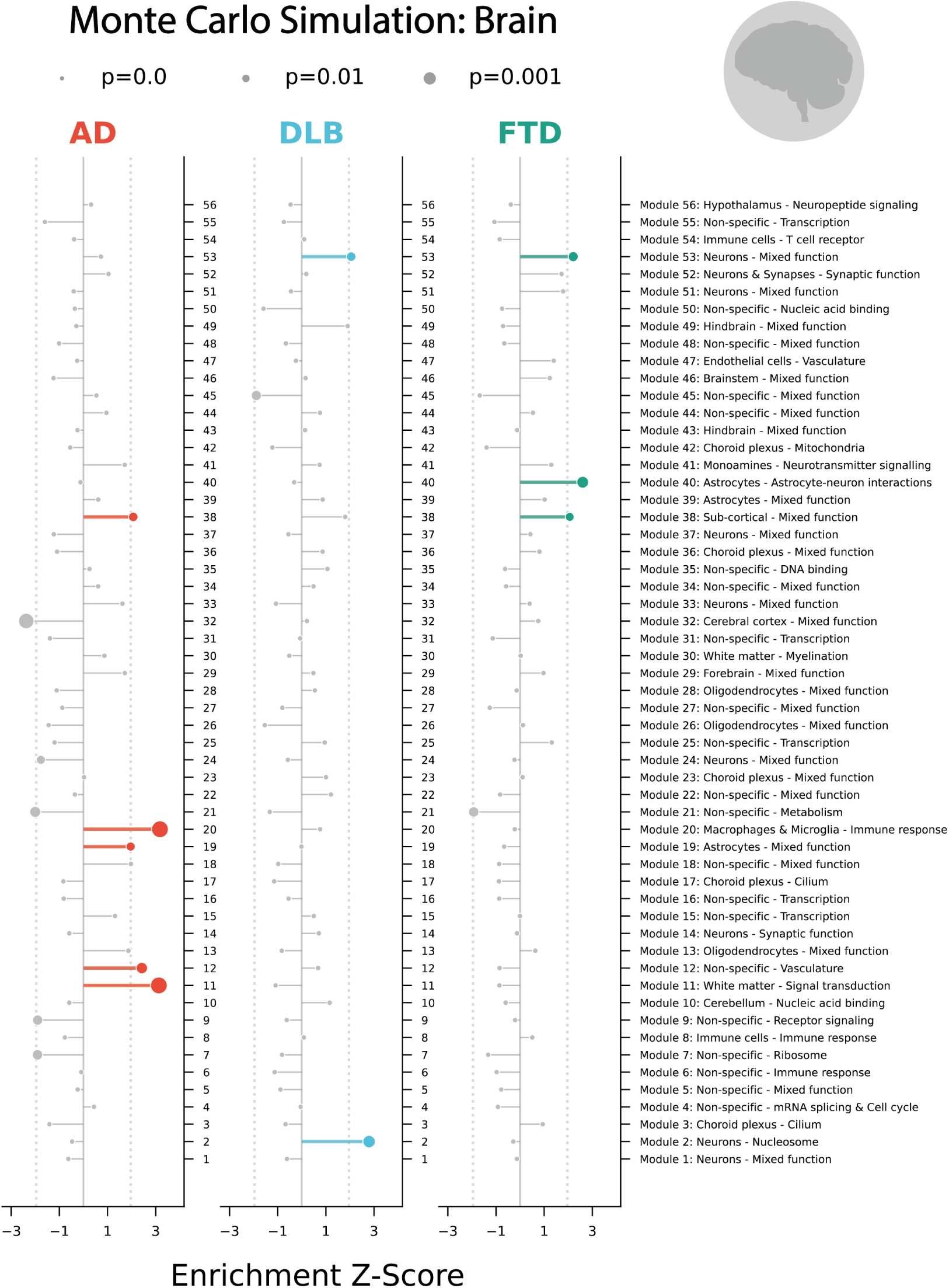
Lollipop plots showing the Monte Carlo simulation disease risk gene enrichment of the brain reigon gene modules per disease. Enrichment of dementia risk genes in brain region gene modules were analyzed through 10^6^ Monte Carlo simulations. All modules enriched *p*_*empirical*_ < 0.05 are highlighted by color, the gray lollipops are not significant or of negative Z-score (AD: Red, DLB: Blue, and FTD: Green). The size of the plot point indicates the p-value (lower *p*_*empirical*_ has a larger dot size).

**Figure 2—figure supplement 3.**
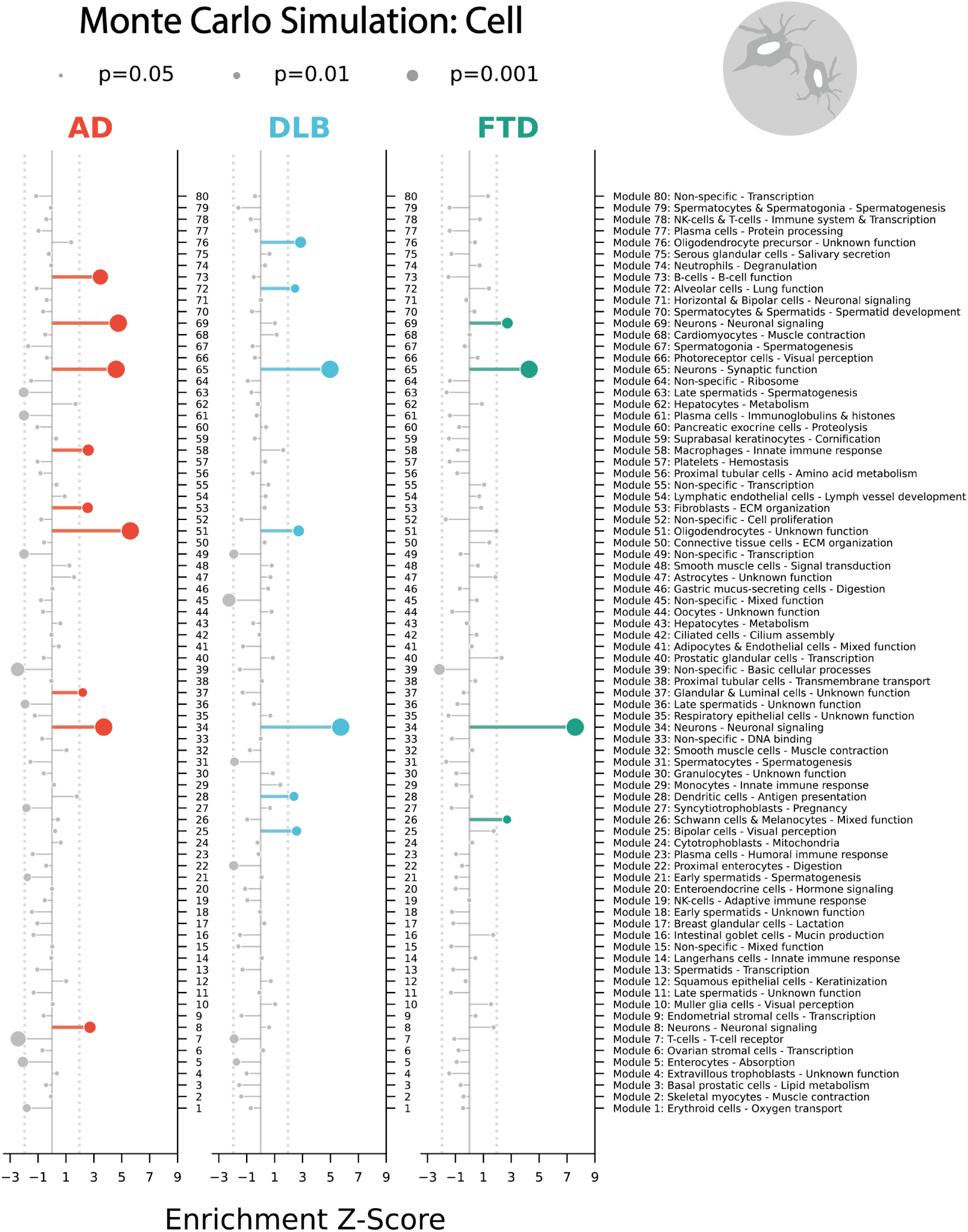
Lollipop plots showing the Monte Carlo simulation disease risk gene enrichment of the cell type gene modules per disease. Enrichment of dementia risk genes in cell gene modules were analyzed through 10^6^ Monte Carlo simulations. All modules enriched *p*_*empirical*_ < 0.05 are highlighted by color, the gray lollipops are not significant or of negative Z-score (AD: Red, DLB: Blue, and FTD: Green). The size of the plot point indicates the p-value (lower *p*_*empirical*_ has a larger dot size).

**Figure 2—figure supplement 4.**
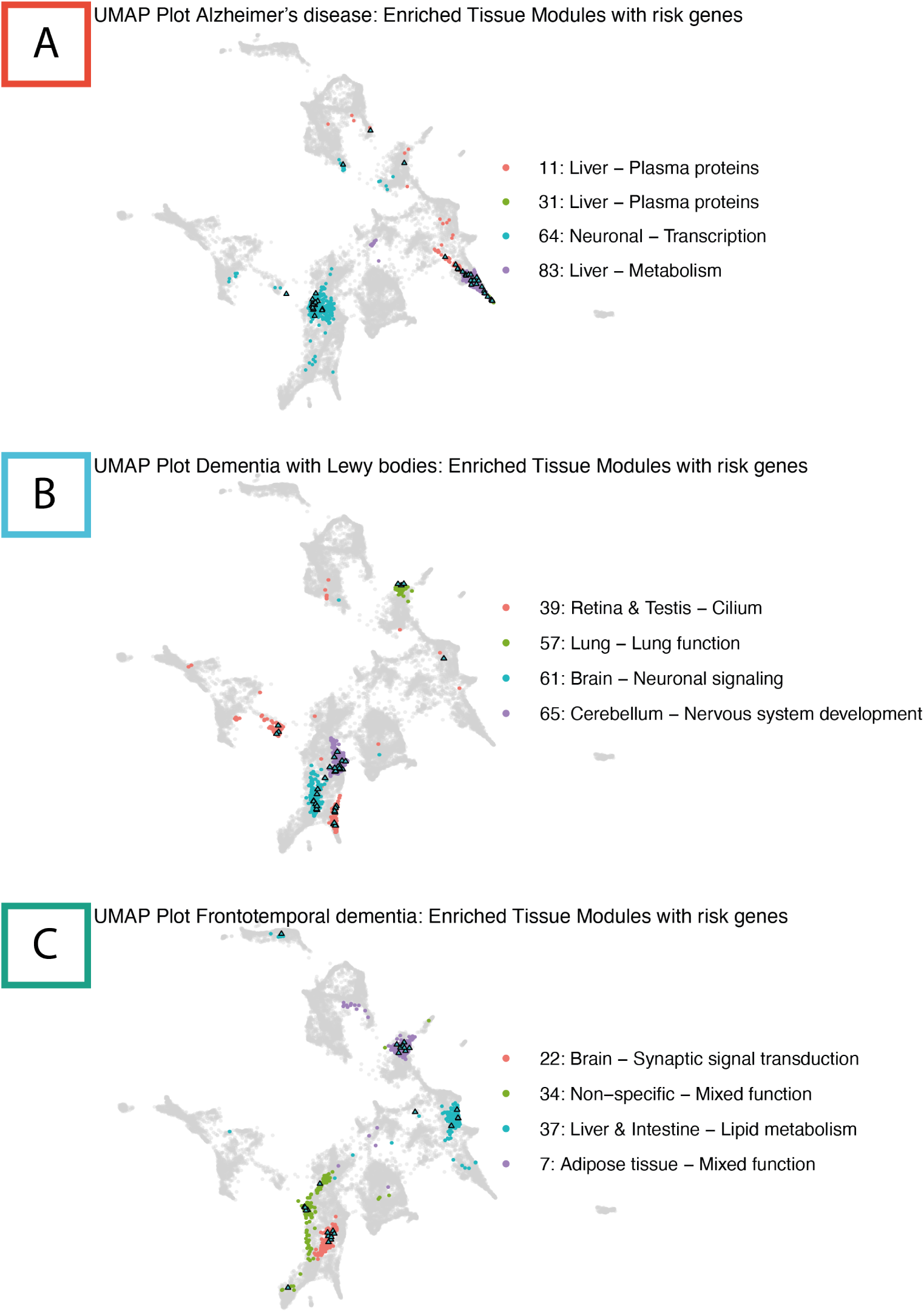
HPA tissue UMAP visualization per disease. UMAP visualization of the gene modules based on the HPA tissue data, highlighting disease gene enriched modules (*p*_*nominal*_ < 0.05). Individual genes are marked by cyan triangle points. The risk genes and enriched modules specific to each disease is depicted in their own UMAP, AD in A; DLB in B; and FTD in C.

**Figure 2—figure supplement 5.**
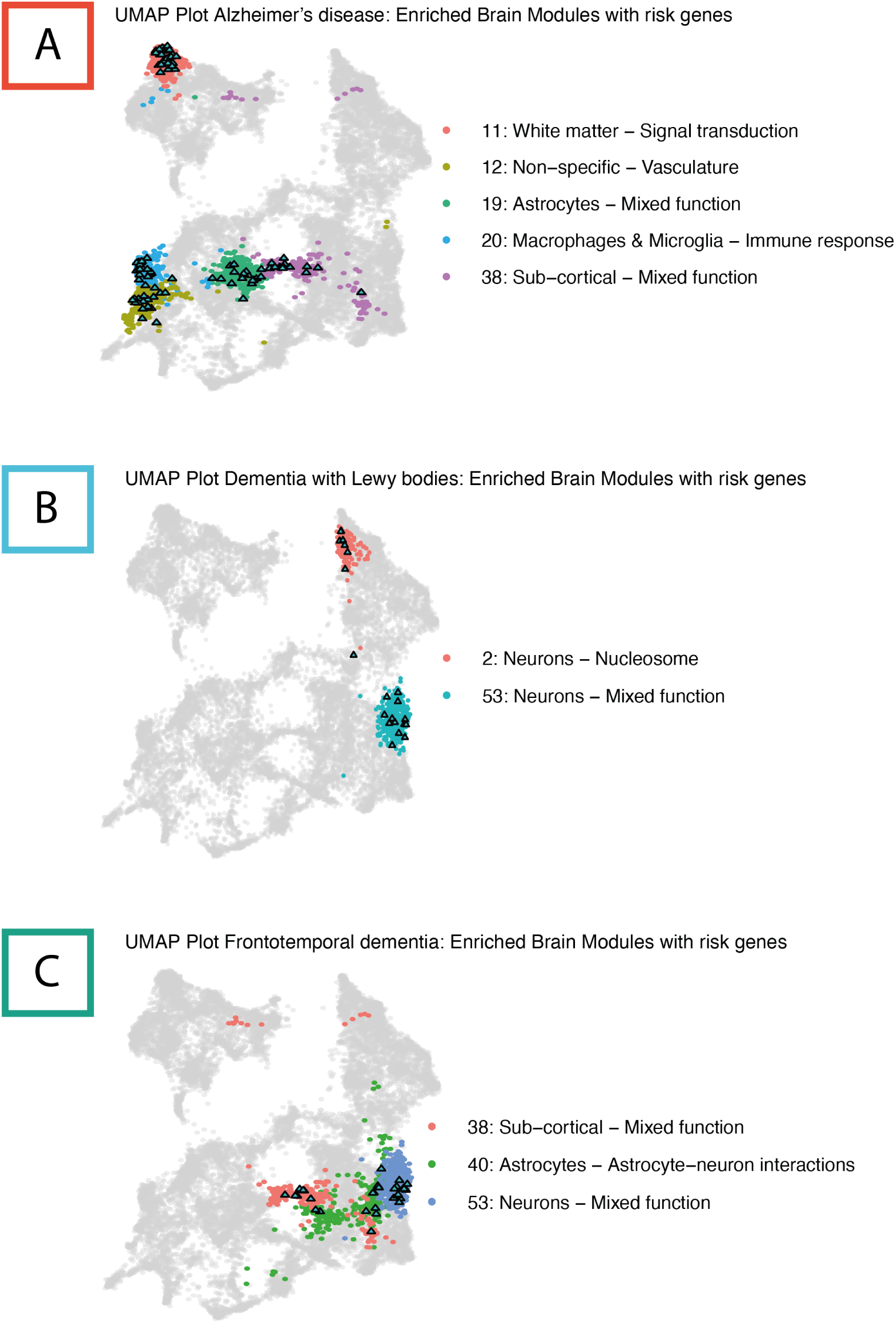
HPA brain region UMAP visualization per disease. UMAP visualization of the gene modules based on the HPA brain data, highlighting disease gene enriched modules (*p*_*nominal*_ < 0.05). Individual genes are marked by cyan triangle points. The risk genes and enriched modules specific to each disease is depicted in their own UMAP, AD in A; DLB in B; and FTD in C.

**Figure 2—figure supplement 6.**
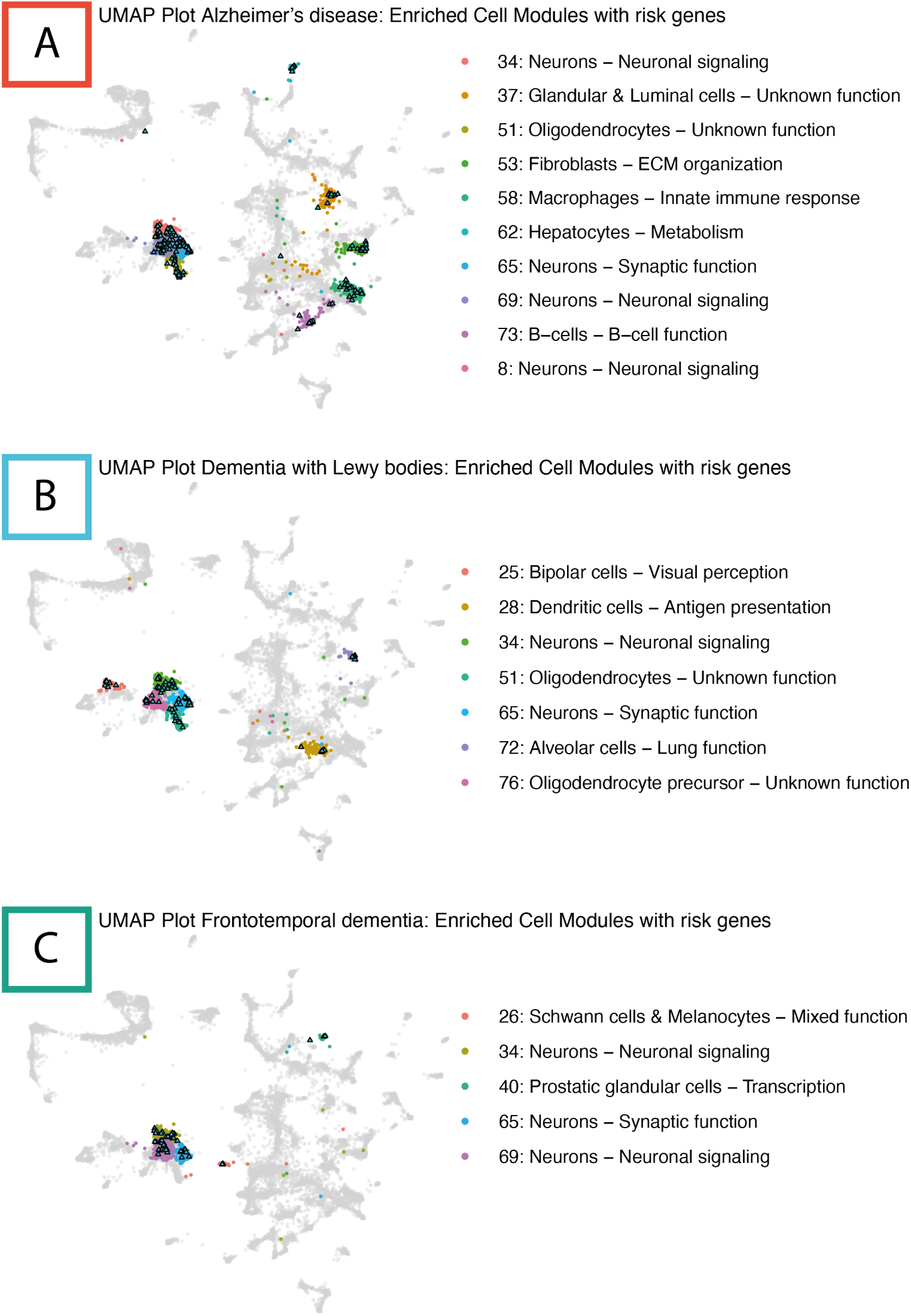
HPA cell type UMAP visualization per disease. UMAP visualization of the gene modules based on the HPA cell data, highlighting disease gene enriched modules (*p*_*nominal*_ < 0.05). Individual genes are marked by cyan triangle points. The risk genes and enriched modules specific to each disease is depicted in their own UMAP, AD in A; DLB in B; and FTD in C.

**Figure 2—figure supplement 7.**
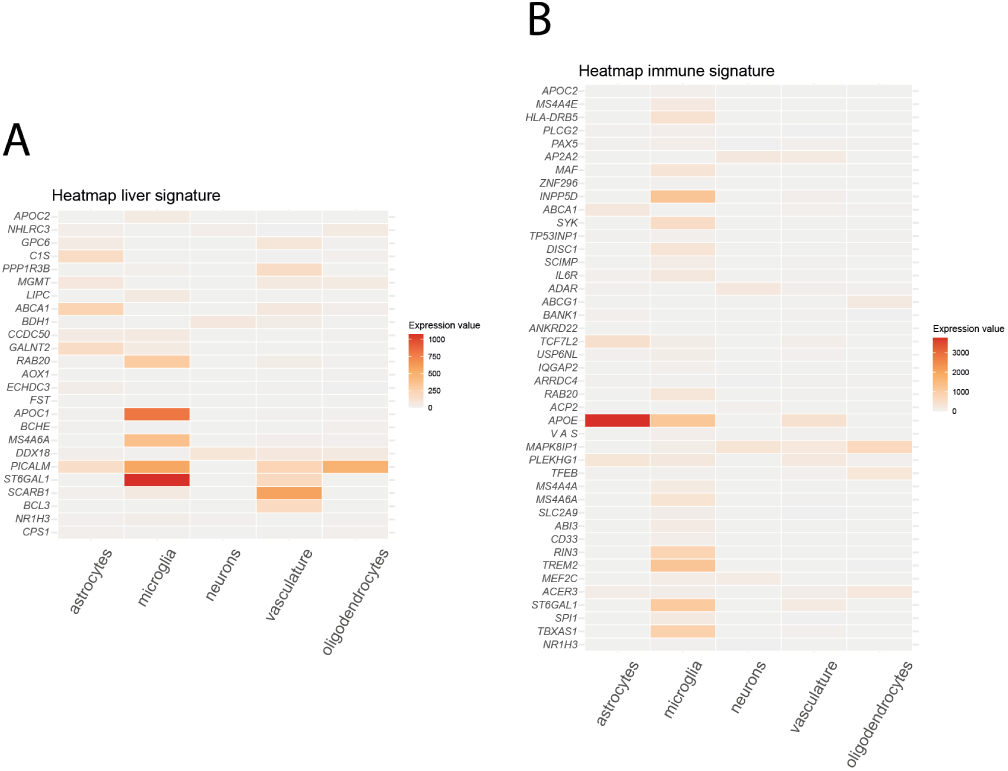
Heatmaps of cell type expression of the liver- and immune associated risk signature genes. Heat map of the Alzheimer’s disease risk genes part of the liver (panel A) and immune (panel B) associated signatures gene expression in spatial sequencing data of HPA human non-demented frontal cortex data. Expression across the cell types; astrocytes, microglia, neurons, vasculature, and oligodendrocytes. Not all genes of the risk signatures were found in the spatial data.

## References

Adori C, Daraio T, Kuiper R, Barde S, Horvathova L, Yoshitake T, Ihnatko R, Valladolid-Acebes I, Vercruysse P, Wellendorf AM, Gramignoli R, Bozoky B, Kehr J, Theodorsson E, Cancelas JA, Mravec B, Jorns C, Ellis E, Mulder J, Uhlén M, et al. Disorganization and degeneration of liver sympathetic innervations in nonalcoholic fatty liver disease revealed by 3D imaging. Sci Adv. 2021; 7:5733–5754.

Aikawa T, Ren Y, Yamazaki Y, Tachibana M, Johnson MR, Anderson CT, Martens YA, Holm ML, Asmann YW, Saito T, Saido TC, Fitzgerald ML, Bu G, Kanekiyo T. ABCA7 haplodeficiency disturbs microglial immune responses in the mouse brain. Proceedings of the National Academy of Sciences of the United States of America. 2019; 116:23790–23796. doi: 10.1073/pnas.1908529116.

Alshehri GH, Al-kuraishy HM, Al-Gareeb AI, Fawzy MN, Waheed HJ, Papadakis M, Alexiou A, Batiha GES. A novel therapeutic prospect: a dual-acting tirzepatide for Alzheimer’s disease. European Journal of Pharmacology. 2025 9; 1003. doi: 10.1016/j.ejphar.2025.177979.

Anita NZ, Zebarth J, Chan B, Wu CY, Syed T, Shahrul D, Nguyen MM, Pakosh M, Herrmann N, Lanctôt KL, Swardfager W. Inflammatory markers in type 2 diabetes with vs. without cognitive impairment; a systematic review and meta-analysis. Brain, Behavior, and Immunity. 2022 2; 100:55–69. doi: 10.1016/j.bbi.2021.11.005.

Ard MD, Cole GM, Wei J, Mehrle AP, Fratkin JD. Scavenging of Alzheimer’s amyloid *β*-protein by microglia in culture. Journal of Neuroscience Research. 1996; 43:190–202. doi: 10.1002/(SICI)1097-4547(19960115)43:2<190::AID-JNR7>3.0.CO;2-B.

Bahia VS, Takada LT, Deramecourt V. Neuropathology of frontotemporal lobar degeneration: a review. Dementia e Neuropsychologia. 2013; 7:19–26. doi: 10.1590/S1980-57642013DN70100004.

Bano D, Ehninger D, Bagetta G. Decoding metabolic signatures in Alzheimer’s disease: a mitochondrial perspective. Cell Death Discovery. 2023 12; 9. doi: 10.1038/s41420-023-01732-3.

Bellenguez C, Küçükali F, Jansen IE, Kleineidam L, Moreno-Grau S, Amin N, Naj AC, Campos-Martin R, Grenier-Boley B, Andrade V, Holmans PA, Boland A, Damotte V, van der Lee SJ, Costa MR, Kuulasmaa T, Yang Q, de Rojas I, Bis JC, Yaqub A, et al. New insights into the genetic etiology of Alzheimer’s disease and related dementias. Nature Genetics. 2022 4; 54:412–436. https://www.nature.com/articles/s41588-022-01024-z, doi: 10.1038/s41588-022-01024-z.

Bianca VD, Dusi S, Bianchini E, Prà ID, Rossi F. *β*-amyloid activates the O2/·- forming NADPH oxidase in microglia, monocytes, and neutrophils. A possible inflammatory mechanism of neuronal damage in Alzheimer’s disease. Journal of Biological Chemistry. 1999 5; 274:15493–15499. doi: 10.1074/jbc.274.22.15493.

Braak H, Thal DR, Ghebremedhin E, Tredici KD. Stages of the pathologic process in alzheimer disease: Age categories from 1 to 100 years. Journal of Neuropathology and Experimental Neurology. 2011; 70:960–969. doi: 10.1097/NEN.0b013e318232a379.

Broce I, Karch CM, Wen N, Fan CC, Wang Y, Tan CH, Kouri N, Ross OA, Höglinger GU, Muller U, Hardy J, Momeni P, Hess CP, Dillon WP, Miller ZA, Bonham LW, Rabinovici GD, Rosen HJ, Schellenberg GD, Franke A, et al. Immune-related genetic enrichment in frontotemporal dementia: An analysis of genome-wide association studies. PLoS Medicine. 2018 1; 15. doi: 10.1371/journal.pmed.1002487.

Brubaker WD, Crane A, Johansson JU, Yen K, Garfinkel K, Mastroeni D, Asok P, Bradt B, Sabbagh M, Wallace TL, Glavis-Bloom C, Tenner AJ, Rogers J. Peripheral complement interactions with amyloid *β* peptide: Erythrocyte clearance mechanism. Alzheimer’s & dementia : the journal of the Alzheimer’s Association. 2017 12; 13:1397. https://pmc.ncbi.nlm.nih.gov/articles/PMC5880643/, doi: 10.1016/J.JALZ.2017.03.010.

Cheon SY, Song J. Novel insights into non-alcoholic fatty liver disease and dementia: insulin resistance, hyper-ammonemia, gut dysbiosis, vascular impairment, and inflammation. Cell and Bioscience. 2022 12; 12. doi: 10.1186/s13578-022-00836-0.

Delvadia P, Dhote V, Mandloi AS, Soni R, Shah J. Dual GLP-1 and GIP Agonist Tirzepatide Exerted Neuroprotective Action in a Parkinson’s Disease Rat Model. ACS Chemical Neuroscience. 2025 3; 16:818–825. doi: 10.1021/acschemneuro.4c00729.

Doens D, Fernández PL. Microglia receptors and their implications in the response to amyloid *β* for Alzheimer’s disease pathogenesis. Journal of Neuroinflammation. 2014 3; 11. doi: 10.1186/1742-2094-11-48.

Espay AJ, Kepp KP, Herrup K. Lecanemab and Donanemab as Therapies for Alzheimer’s Disease: An Illustrated Perspective on the Data. eNeuro. 2024 7; 11. doi: 10.1523/ENEURO.0319-23.2024.

Estrada LD, Ahumada P, Cabrera D, Arab JP. Liver Dysfunction as a Novel Player in Alzheimer’s Progression: Looking Outside the Brain. Frontiers in Aging Neuroscience. 2019 7; 11. doi: 10.3389/fnagi.2019.00174.

Fanning S, Selkoe D, Dettmer U. Parkinson’s disease: proteinopathy or lipidopathy? npj Parkinson’s Disease. 2020 12; 6. doi: 10.1038/s41531-019-0103-7.

Ferrari R, Hernandez DG, Nalls MA, Rohrer JD, Ramasamy A, Kwok JBJ, Dobson-Stone C, S BSBW, Schofield PR, Halliday GM, Hodges JR, Piguet O, Bartley L, Thompson E, Haan E, Hernández I, Ruiz A, Boada M, Borroni B, Padovani A, et al. Frontotemporal dementia and its subtypes: A genome-wide association study. The Lancet Neurology. 2014; 13:686–699. doi: 10.1016/S1474-4422(14)70065-1.

Filho FLW, Lopes PRM, de Almeida AM, Sano VKT, Tamashiro FM, Gonçalves OR, de Moraes FCA, Kreuz M, Kelly FA, Feitoza PVS. Statin use and dementia risk: A systematic review and updated meta-analysis. Alzheimer’s and Dementia: Translational Research and Clinical Interventions. 2025 1; 11. doi: 10.1002/trc2.70039.

Filipović B, Marković O, Urić V, Filipović B. Cognitive Changes and Brain Volume Reduction in Patients with Nonalcoholic Fatty Liver Disease. Canadian Journal of Gastroenterology and Hepatology. 2018; 2018. doi: 10.1155/2018/9638797.

Gerrits E, Brouwer N, Kooistra SM, Woodbury ME, Vermeiren Y, Lambourne M, Mulder J, Kummer M, Möller T, Biber K, den Dunnen WFA, Deyn PPD, Eggen BJL, Boddeke EWGM. Distinct amyloid-*β* and tau-associated microglia profiles in Alzheimer’s disease. Acta Neuropathologica. 2021 5; 141:681–696. doi: 10.1007/s00401-021-02263-w.

Iqbal A, Baldrighi M, Murdoch JN, Fleming A, Wilkinson CJ. Alpha-synuclein aggresomes inhibit ciliogenesis and multiple functions of the centrosome. Biology Open. 2020 10; 9. doi: 10.1242/bio.054338.

Jay TR, Hirsch AM, Broihier ML, Miller CM, Neilson LE, Ransohoff RM, Lamb BT, Landreth GE. Disease progression-dependent effects of TREM2 deficiency in a mouse model of Alzheimer’s disease. Journal of Neuroscience. 2017 1; 37:637–647. doi: 10.1523/JNEUROSCI.2110-16.2016.

Kanekiyo T, Bu G. The low-density lipoprotein receptor-related protein 1 and amyloid-*β* clearance in Alzheimer’s disease. Frontiers in Aging Neuroscience. 2014; 6. doi: 10.3389/fnagi.2014.00093.

Kardinal R, Wachten D. Macrophages in metaflammation – fueling chronic inflammation in metabolic disease. Pflugers Archiv European Journal of Physiology. 2026 1; 478. doi: 10.1007/s00424-025-03141-0.

Karlsson M, Zhang C, Méar L, Zhong W, Digre A, Katona B, Sjöstedt E, Butler L, Odeberg J, Dusart P, Edfors F, Oksvold P, von Feilitzen K, Zwahlen M, Arif M, Altay O, Li X, Ozcan M, Mardinoglu A, Fagerberg L, et al. A single-cell type transcriptomics map of human tissues. Sci Adv. 2021; 7. www.proteinatlas.org.

Khan SS, Jaimon E, Lin YE, Nikoloff J, Tonelli F, Alessi DR, Pfeffer SR. Loss of primary cilia and dopaminergic neuroprotection in pathogenic LRRK2-driven and idiopathic Parkinson’s disease. Proceedings of the National Academy of Sciences of the United States of America. 2024 8; 121. doi: 10.1073/pnas.2402206121.

Knox C, Wilson M, Klinger CM, Franklin M, Oler E, Wilson A, Pon A, Cox J, Chin NEL, Strawbridge SA, Garcia-Patino M, Kruger R, Sivakumaran A, Sanford S, Doshi R, Khetarpal N, Fatokun O, Doucet D, Zubkowski A, Rayat DY, et al. DrugBank 6.0: the DrugBank Knowledgebase for 2024. Nucleic Acids Research. 2024 1; 52:D1265–D1275. doi: 10.1093/nar/gkad976.

Kunkle BW, Grenier-Boley B, Sims R, Bis JC, Damotte V, Naj AC, Boland A, Vronskaya M, van der Lee SJ, Amlie-Wolf A, Bellenguez C, Frizatti A, Chouraki V, Martin ER, Sleegers K, Badarinarayan N, Jakobsdottir J, Hamilton-Nelson KL, Moreno-Grau S, Olaso R, et al. Genetic meta-analysis of diagnosed Alzheimer’s disease identifies new risk loci and implicates A*β*, tau, immunity and lipid processing. Nature Genetics. 2019 3; 51:414–430. https://www.nature.com/articles/s41588-019-0358-2, doi: 10.1038/s41588-019-0358-2.

Livingston G, Huntley J, Liu KY, Costafreda SG, Selbæk G, Alladi S, Ames D, Banerjee S, Burns A, Brayne C, Fox NC, Ferri CP, Gitlin LN, Howard R, Kales HC, Kivimäki M, Larson EB, Nakasujja N, Rockwood K, Samus Q, et al. Dementia prevention, intervention, and care: 2024 report of the Lancet standing Commission. The Lancet. 2024 8; 404:572–628. doi: 10.1016/S0140-6736(24)01296-0.

Lu Y, Pike JR, Hoogeveen RC, Walker KA, Raffield LM, Selvin E, Avery CL, Engel SM, Mielke MM, Garcia T, Palta P. Liver integrity and the risk of Alzheimer’s disease and related dementias. Alzheimer’s and Dementia. 2024 3; 20:1913–1922. doi: 10.1002/ALZ.13601.

Lue LF, Walker DG, Brachova L, Beach TG, Rogers J, Schmidt AM, Stern DM, Yan SD. Involvement of microglial receptor for advanced glycation endproducts (RAGE)in Alzheimer’s disease: Identification of a cellular activation mechanism. Experimental Neurology. 2001; 171:29–45. doi: 10.1006/exnr.2001.7732.

Magno R, Maia AT. Gwasrapidd: An R package to query, download and wrangle GWAS catalog data. Bioinfor-matics. 2020 1; 36:649–650. doi: 10.1093/bioinformatics/btz605.

Matošević A, Opsenica DM, Bartolić M, Maraković N, Stoilković A, Komatović K, Zandona A, Žunec S, Bosak A. Derivatives of Amodiaquine as Potent Human Cholinesterases Inhibitors: Implication for Treatment of Alzheimer’s Disease. Molecules. 2024 11; 29. doi: 10.3390/molecules29225357.

Mishra A, Ferrari R, Heutink P, Hardy J, Pijnenburg Y, Posthuma D. Gene-based association studies report genetic links for clinical subtypes of frontotemporal dementia. Brain. 2017 5; 140:1437–1446. doi: 10.1093/brain/awx066.

Monte SMDL, Wands JR. Review of insulin and insulin-like growth factor expression, signaling, and malfunction in the central nervous system: Relevance to Alzheimer’s disease. Journal of Alzheimer’s Disease. 2005; 7:45–61.

Orme T, Guerreiro R, Bras J. The Genetics of Dementia with Lewy Bodies: Current Understanding and Future Directions. Current Neurology and Neuroscience Reports. 2018 10; 18. doi: 10.1007/s11910-018-0874-y.

Park SJ, Diaz JG, Um E, Hahn YS. Major roles of kupffer cells and macrophages in NAFLD development. Frontiers in Endocrinology. 2023; 14. doi: 10.3389/fendo.2023.1150118.

Raffaele F, Claudia M, John H. Genetics and molecular mechanisms of frontotemporal lobar degeneration: an update and future avenues. Neurobiology of Aging. 2019 6; 78:98–110. doi: 10.1016/j.neurobiolaging.2019.02.006.

Reitz C, Brayne C, Mayeux R. Epidemiology of Alzheimer disease. Nature Reviews Neurology. 2011 3; 7:137–152. doi: 10.1038/nrneurol.2011.2.

Rohrer JD, Guerreiro MR, Vandrovcova J, Uphill J, Reiman BD, Beck J, Isaacs BAM, Authier DA, Ferrari MR, Fox NC, Mackenzie IRA, Warren JD, Silva FRD, Holton DJ, Revesz T, Hardy J, Mead S, Rossor MN. The heritability and genetics of frontotemporal lobar degeneration. Neurology. 2009; doi: 10.1212/WNL.0b013e3181bf997a.

de Ruyter FJH, Morrema THJ, den Haan J, Gase G, Twisk JWR, de Boer JF, Scheltens P, Bouwman FH, Verbraak FD, Rozemuller AJM, Hoozemans JJM. *α*-Synuclein pathology in post-mortem retina and optic nerve is specific for *α*-synucleinopathies. npj Parkinson’s Disease. 2023 12; 9. doi: 10.1038/s41531-023-00570-5.

Sanghvi H, Singh R, Morrin H, Rajkumar AP. Systematic review of genetic association studies in people with Lewy body dementia. International Journal of Geriatric Psychiatry. 2020 5; 35:436–448. doi: 10.1002/gps.5260.

Schmidt S, Luecken MD, Trümbach D, Hembach S, Niedermeier KM, Wenck N, Pflügler K, Stautner C, Böttcher A, Lickert H, Ramirez-Suastegui C, Ahmad R, Ziller MJ, Fitzgerald JC, Ruf V, van de Berg WDJ, Jonker AJ, Gasser T, Winner B, Winkler J, et al. Primary cilia and SHH signaling impairments in human and mouse models of Parkinson’s disease. Nature Communications. 2022 12; 13. doi: 10.1038/s41467-022-32229-9.

Serpieri V, D’Abrusco F, Valente EM. The relevance of primary cilia in neurological disorders. The Lancet Neurology. 2025 9; 24:763–775. doi: 10.1016/S1474-4422(25)00226-1.

Sherva R, Bayly H, Zhang R, Mez J, Hauger RL, Merritt VC, Panizzon MS, Gaziano JM, Farrer LA, Logue MW. GWAS of 205,500 Alzheimer’s disease and related dementia cases reveals 19 novel, European specific and 11 cross-ancestry risk loci. Alzheimer’s & Dementia. 2025 12; 21. https://alz-journals.onlinelibrary.wiley.com/doi/10.1002/alz70855_105500, doi: 10.1002/alz70855_105500.

Sjöstedt E, Zhong W, Fagerberg L, Karlsson M, Mitsios N, Adori C, Oksvold P, Edfors F, Limiszewska A, Hikmet F, Huang J, Du Y, Lin L, Dong Z, Yang L, Liu X, Jiang H, Xu X, Wang J, Yang H, et al. An atlas of the protein-coding genes in the human, pig, and mouse brain. Science. 2020 3; 367. doi: 10.1126/science.aay5947.

Song D, Li Y, Yang LL, Luo YX, Yao XQ. Bridging systemic metabolic dysfunction and Alzheimer’s disease: the liver interface. Molecular Neurodegeneration. 2025 12; 20. doi: 10.1186/s13024-025-00849-6.

Steen E, Terry BM, Rivera EJ, Cannon JL, Neely TR, Tavares RT, Xu XJ, Wands JR, de la Mont SM. Impaired insulin and insulin-like growth factor expression and signaling mechanisms in Alzheimer’s disease – is this type 3 diabetes? Journal of Alzheimer’s Disease. 2005 2; 7:63–80. doi: 10.3233/jad-2005-7107.

Sudwarts A, Ramesha S, Gao T, Ponnusamy M, Wang S, Hansen M, Kozlova A, Bitarafan S, Kumar P, Beaulieu-Abdelahad D, Zhang X, Collier L, Szekeres C, Wood LB, Duan J, Thinakaran G, Rangaraju S. BIN1 is a key regulator of proinflammatory and neurodegeneration-related activation in microglia. Molecular Neurodegeneration. 2022 12; 17. doi: 10.1186/s13024-022-00535-x.

Tian Z, Zhang Y, Xu J, Yang Q, Hu D, Feng J, Gai C. Primary cilia in Parkinson’s disease: summative roles in signaling pathways, genes, defective mitochondrial function, and substantia nigra dopaminergic neurons. Frontiers in Aging Neuroscience. 2024; 16. doi: 10.3389/fnagi.2024.1451655.

Uhlén M, Fagerberg L, Hallström BM, Lindskog C, Oksvold P, Mardinoglu A, Åsa Sivertsson, Kampf C, Sjöstedt E, Asplund A, Olsson IM, Edlund K, Lundberg E, Navani S, Szigyarto CAK, Odeberg J, Djureinovic D, Takanen JO, Hober S, Alm T, et al. Tissue-based map of the human proteome. Science. 2015 1; 347. doi: 10.1126/sci-ence.1260419.

Varma VR, Desai RJ, Navakkode S, Wong LW, Anerillas C, Loeffler T, Schilcher I, Mahesri M, Chin K, Horton DB, Kim SC, Gerhard T, Segal JB, Schneeweiss S, Gorospe M, Sajikumar S, Thambisetty M. Hydroxychloroquine lowers Alzheimer’s disease and related dementias risk and rescues molecular phenotypes related to Alzheimer’s disease. Molecular Psychiatry. 2023 3; 28:1312–1326. doi: 10.1038/s41380-022-01912-0.

Wang Y, Cella M, Mallinson K, Ulrich JD, Young KL, Robinette ML, Gilfillan S, Krishnan GM, Sudhakar S, Zin-selmeyer BH, Holtzman DM, Cirrito JR, Colonna M. TREM2 lipid sensing sustains the microglial response in an Alzheimer’s disease model. Cell. 2015 3; 160:1061–1071. doi: 10.1016/j.cell.2015.01.049.

Weinshenker D. Long Road to Ruin: Noradrenergic Dysfunction in Neurodegenerative Disease. Trends in Neu-rosciences. 2018; 41:211–223. 10.1016/j.tins.2018.01.010, doi: 10.1016/j.tins.2018.01.010.

Weinstein G, O’Donnell A, Davis-Plourde K, Zelber-Sagi S, Ghosh S, Decarli CS, Thibault EG, Sperling RA, Johnson KA, Beiser AS, Seshadri S. Non-Alcoholic Fatty Liver Disease, Liver Fibrosis, and Regional Amyloid-*β* and Tau Pathology in Middle-Aged Adults: The Framingham Study. Journal of Alzheimer’s Disease. 2022; 86:1371–1383. doi: 10.3233/JAD-215409.

Wensel TG, Potter VL, Moye A, Zhang Z, Robichaux MA. Structure and dynamics of photoreceptor sensory cilia. Pflugers Archiv European Journal of Physiology. 2021 9; 473:1517–1537. doi: 10.1007/s00424-021-02564-9.

Wilson DM, Cookson MR, Bosch LVD, Zetterberg H, Holtzman DM, Dewachter I. Hallmarks of neurodegenerative diseases. Cell. 2023 2; 186:693–714. doi: 10.1016/j.cell.2022.12.032.

Wirdefeldt K, Gatz M, Reynolds CA, Prescott CA, Pedersen NL. Heritability of Parkinson disease in Swedish twins: A longitudinal study. Neurobiology of Aging. 2011; 32:1923.e1–1923.e8. doi: 10.1016/j.neurobiolaging.2011.02.017.

Xie Z, Harris-White ME, Wals PA, Frautschy SA, Finch CE, Morgan TE. Apolipoprotein J (clusterin) activates rodent microglia in vivo and in vitro. Journal of Neurochemistry. 2005 5; 93:1038–1046. doi: 10.1111/j.1471-4159.2005.03065.x.

Yang S, Zhao X, Zhang Y, Tang Q, Li Y, Du Y, yu P. Tirzepatide shows neuroprotective effects via regulating brain glucose metabolism in APP/PS1 mice. Peptides. 2024 9; 179. doi: 10.1016/j.peptides.2024.171271.

Yuan S, Wang Y, Yang J, Tang Y, Wu W, Meng X, Jian Y, Lei Y, Liu Y, Tang C, Zhao Z, Zhao F, Liu W. Treadmill exercise can regulate the redox balance in the livers of APP/PS1 mice and reduce LPS accumulation in their brains through the gut-liver-kupffer cell axis. Aging (Albany NY). 2024; 16:1374. https://pmc.ncbi.nlm.nih.gov/articles/PMC10866404/, doi: 10.18632/AGING.205432.

Zarghamravanbakhsh P, Frenkel M, Poretsky L. Metabolic causes and consequences of nonalcoholic fatty liver disease (NAFLD). Metabolism Open. 2021 12; 12:100149. doi: 10.1016/j.metop.2021.100149.

Zhang J, Wang Y, Zhang Y, Yao J. Genome-wide association study in Alzheimer’s disease: a bibliometric and visualization analysis. Frontiers in Aging Neuroscience. 2023; 15. doi: 10.3389/fnagi.2023.1290657.

